# Hopanoid lipids promote soybean-*Bradyrhizobium* symbiosis

**DOI:** 10.1101/2023.09.04.556284

**Authors:** Huiqiao Pan, Ashley Shim, Matthew B. Lubin, Brittany J. Belin

**Affiliations:** Department of Embryology, Carnegie Institution for Science, Baltimore, MD 21218, United States; Department of Biology, Johns Hopkins University, Baltimore, MD 21218, United States

**Keywords:** hopanoids, lipids, symbiosis, *Bradyrhizobium*, soybean, *Aeschynomene*, root nodules, nitrogen fixation, flagella, secretion system

## Abstract

The symbioses between leguminous plants and nitrogen-fixing bacteria known as rhizobia are well known for promoting plant growth and sustainably increasing soil nitrogen. Recent evidence indicates that hopanoids, a family of steroid-like lipids, promote *Bradyrhizobium* symbioses with tropical legumes. To characterize hopanoids in *Bradyrhizobium* symbiosis with soybean, the most economically significant *Bradyrhizobium* host, we validated a recently published cumate-inducible hopanoid mutant of *Bradyrhizobium diazoefficiens* USDA110, Pcu-*shc*::Δ*shc*. GC-MS analysis showed that this strain does not produce hopanoids without cumate induction, and under this condition, is impaired in growth in rich medium and under osmotic, temperature, and pH stress. *In planta*, Pcu-*shc*::Δ*shc* is an inefficient soybean symbiont with significantly lower rates of nitrogen fixation and low survival within host tissue. RNA-seq revealed that hopanoid loss reduces expression of flagellar motility and chemotaxis-related genes, further confirmed by swim plate assays, and enhances expression of genes related to nitrogen metabolism and protein secretion. These results suggest that hopanoids provide a significant fitness advantage to *B. diazoefficiens* in legume hosts and provide a foundation for future mechanistic studies of hopanoid function in protein secretion and motility.

**IMPORTANCE:** A major problem for global sustainability is feeding our exponentially growing human population while available arable land is decreasing, especially in areas with the greatest population growth. Harnessing the power of plant-beneficial microbes has gained attention as a potential solution, including the increasing our reliance on the symbioses of leguminous plants and nitrogen-fixing rhizobia. This study examines the role of hopanoid lipids in the symbiosis between *Bradyrhizobium diazoefficiens* USDA110, an important commercial inoculant strain, and its economically important host soybean. Our research extends our knowledge of the functions of bacterial lipids in symbiosis to an agricultural context, which may one day help improve the practical applications of plant-beneficial microbes in agriculture.

## INTRODUCTION

Nitrogen-fixing soil bacteria known as rhizobia are major contributors to the global nitrogen cycle and engage in economically- and environmentally-significant symbioses with legume plants (1). These symbioses help reduce the amount of nitrogen fertilizers required in commercial agriculture, ultimately lowering agricultural greenhouse gas emissions (2). Understanding the rhizobial genes that promote these symbioses has the potential to enhance agricultural sustainability.

In symbioses between legumes and the *Bradyrhizobium* genus of rhizobia, previous work has demonstrated that efficient symbiosis requires bacterial hopanoid lipids (3–5). Hopanoids are a family of pentacyclic lipids that are produced by diverse bacteria (6–18) and are strongly conserved in nitrogen-fixing symbionts of plants (12, 19). First studied as biomarkers of ancient bacteria (20, 21), hopanoids are bacterial analogs of cholesterol that can modify cell membrane biophysics (5, 22–26) and promote tolerance of stress from antibiotics, detergents, pH, elevated temperature, and osmotic pressure (3, 9–11, 24, 27).

In the symbiotic context, prior work on hopanoids has focused on the interactions between *Bradyrhizobia* and the *Aeschynomene* genus of semi-aquatic plants, a broadly distributed forage legume in the developing world. In the photosynthetic *Bradyrhizobium* BTAi1, a deletion strain of the *shc* gene encoding squalene-hopene cyclase, the enzyme that catalyzes the first committed step of hopanoid biosynthesis (**Fig. 1A**), was found to have lower survival and aberrant bacteroid morphology in its native *Aeschynomene evenia* host (5). In *Bradyrhizobium diazoefficiens* USDA110, a native soybean (*Glycine max*) symbiont with a broad range of compatible hosts, only mutant strains that lack specific hopanoid subclasses have been examined. *B. diazoefficiens* strain lacking the hopanoid 2-methylase HpnP has no phenotype in symbiosis with *Aeschynomene afraspera* or soybean, whereas loss of the enzyme HpnH that generates hopanoids with extended hydrocarbon tails inhibits symbiosis with *A. afraspera* but not soybean (3). Further analysis found that *hpnH* deletion mutant slows the development of the symbiosis-specific root nodule organ in *A. afraspera*, in part due to the strain’s reduced motility and poor *in planta* survival (4).

**Figure 1.**
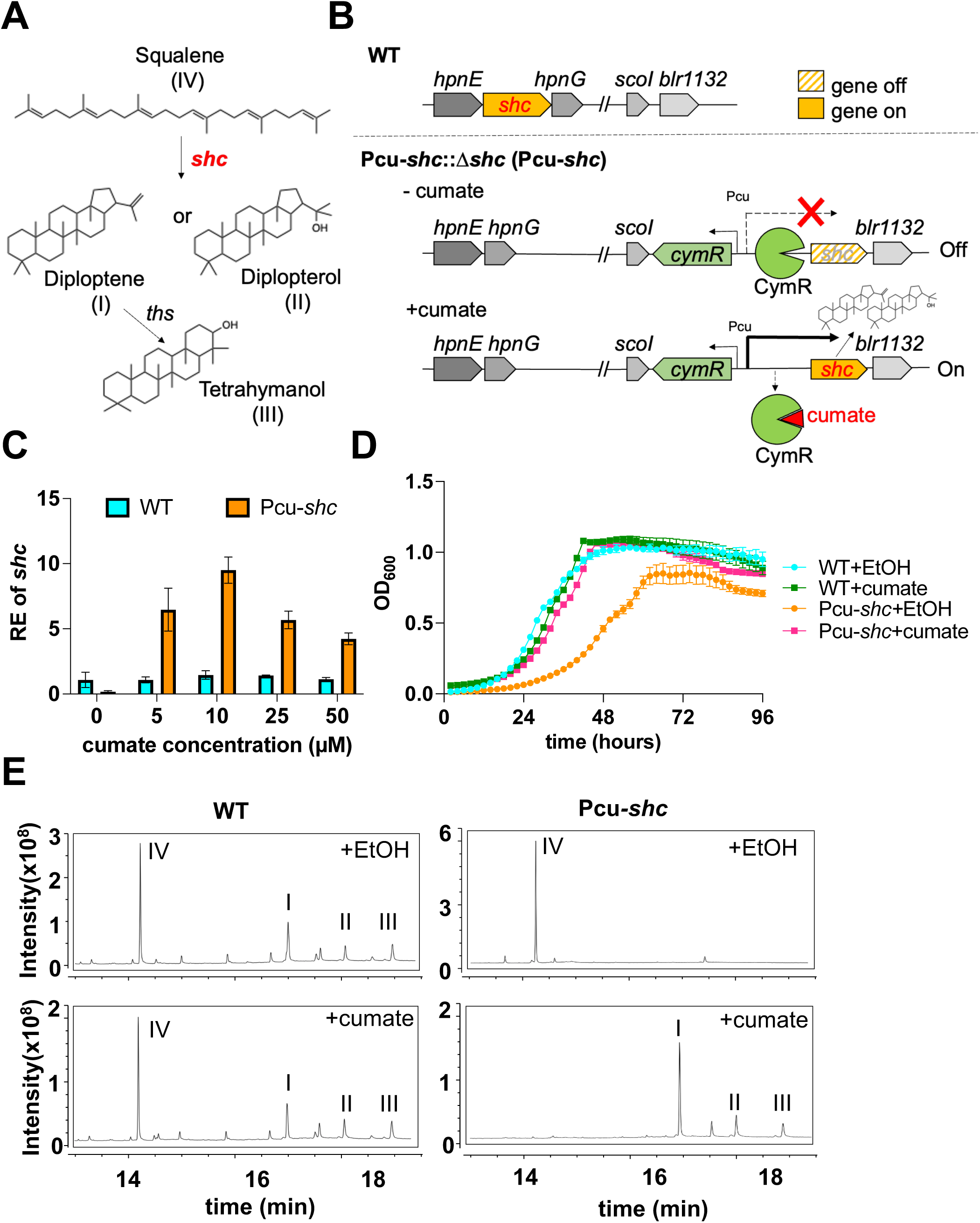
Validation of a cumate-inducible shc strain Pcu-*shc*::Δ*shc* (Pcu-*shc*) of B. *diazoefficiens* USDA 110. (A) The first committed step of hopanoid biosynthesis: squalene-hopene cyclase encoded by *shc* (*hpnF*, blr3004), cyclizes linear squalene (IV) into C_30_ short hopanoids, diploptene (I) and/or diplopterol (II). The tetrahymanol synthase encoded by *ths* (blr0371) converts diploptene to tetrahymanol (III) (41). **(B)** Schematic of WT *B. diazoefficiens* and the cumate-inducible Pcu-*shc* strain. The exogenous cumate-inducible system was integrated into the genomic region between *scoI* (*blr1131*) and *blr1132* in a background of *shc* deletion. This system includes a copy of *shc* driven by a cumate-inducible promoter (Pcu) and a transcriptional repressor gene *cymR* driven by a constitutive promoter. **(C)** qRT-PCR of *shc* in WT and Pcu-*shc* in early exponential phase (OD_600_=0.5-0.8) with a variety of cumate concentrations. The internal control gene was 16S rRNA and relative expression (RE) of *shc* was calculated by ΔΔCt method. **(D)** 4 day growth curves (OD_600_) of WT and Pcu-*shc* with (WT+cumate and Pcu-*shc*+cumate) and without (WT+EtOH and Pcu-*shc*+EtOH) 25 µM cumate in AG media at pH 6.6, 30°C. WT+EtOH and Pcu-*shc*+EtOH (abbr. WT+E and Pcu-*shc*+E hereafter) indicate that the strains were supplemented with an aliquot of ethanol which was used as a solvent of cumate as WT+cumate and Pcu-shc+cumate (abbr. WT+c andPcu-*shc*+c hereafter). **(E)** Total ion chromatograms of lipids extracted from WT and Pcu-*shc* at early exponential phase by gas chromatography–mass spectrometry (GC-MS). Roman numerals represent the same compounds as in (A).

It is not known whether these hopanoid mutant phenotypes are similar to wild type in soybean hosts because hopanoids are not as relevant in soybean as in *Aeschynomene*, or whether the knockout of only certain hopanoid subclasses is insufficient. The roles of hopanoids in *Bradyrhizobium*-soybean symbiosis have been difficult to study, as the *shc* gene required for hopanoid synthesis appears to be essential in the main soybean symbiont, *Bradyrhizobium diazoefficiens* USDA110 (3, 28). Answering this question is an important step in determining the relevance of hopanoids for agriculture, given the global dominance of soybeans as cash crops.

In this study, we examine the effects of complete hopanoid loss on the symbioses between *B. diazoefficiens* and soybean using a recently published cumate-inducible *shc* knockout strain (24). We find that loss of *shc* expression is sufficient to deplete membrane hopanoids and results in similar, though more severe, stress sensitivity phenotypes as observed for other hopanoid mutants. Total hopanoid depletion also strongly inhibits *B. diazoefficiens*-soybean symbiosis over an extended period, with reduced nodule biomass and *in planta* bacterial survival. In contrast, complete hopanoids loss prevents *B. diazoefficiens* symbiosis with *A. afraspera*. Subsequent RNA-seq analyses suggest that these symbiotic phenotypes may result from upregulation of protein secretion and nitrogen assimilation and the disruption of LPS biosynthesis and flagellar motility, the latter of which is confirmed by loss of swimming motility in the *shc* depletion conditions. Together, this work validates a new tool for exploring hopanoid function in *B. diazoefficiens*, expands the relevance of hopanoids in symbiosis to a major crop plant, and suggests potential molecular mechanisms for hopanoids in legume-rhizobia symbiosis.

## RESULTS

### Validation of a cumate-inducible *shc* mutant strain of *B.diazoefficiens*

To characterize hopanoid functions in *B. diazoefficiens*, a cumate-inducible *shc* mutant strain Pcu-*shc*::Δ*shc*, abbreviated as Pcu-*shc* hereafter, was constructed as previously published (24). Briefly, an exogenous copy of *shc* was placed after a cumate-inducible promoter (Pcu) and integrated into the genome, after which the endogenous *shc* was deleted (**Fig. 1B).** We first attempted to validate this strain by quantifying the dose-effect of cumate on *shc* gene expression and bacterial growth. We performed qRT-PCR (ΔΔCt method with an internal control of 16S rRNA) to measure the relative expression (RE) of *shc* in WT and Pcu-*shc* strains after treatment with 5-50 µM cumate or an ethanol (EtOH) vehicle control (**Fig. 1C**). We found that expression of *shc* was insensitive to cumate in WT, whereas the RE of *shc* in Pcu-*shc* was approximately 10-fold lower than WT in the absence of cumate. The mutant strain had 5-10-fold higher *shc* expression than WT after cumate treatment, though we saw minimal changes in *shc* expression within the range of cumate concentrations tested. This result confirmed transcriptional induction of *shc* in Pcu-*shc* by cumate.

In parallel, we performed growth curves of the two strains in rich medium (AG) on a plate reader for samples treated with either 25µM cumate or an EtOH control. WT grown with cumate had similar growth dynamics to WT with EtOH, suggesting that at 25µM concentrations, cumate does not affect WT growth. In contrast, Pcu-*shc* without cumate had pronounced growth defects in both exponential and stationary phases compared to WT+EtOH, and growth was restored to WT levels with 25 µM cumate supplementation **(Fig. 1D)**. We also found that high doses of cumate inhibited bacterial growth **(Fig. S1)**, and therefore chose 25 µM cumate for all following assays, consistent with previous studies (24).

To determine if the low levels of *shc* expression in Pcu-*shc* without cumate are sufficient to deplete hopanoids from the membrane, we used GC-MS (gas chromatography–mass spectrometry) protocols adapted from previous studies (27) to extract and analyze short hopanoids from *B. diazoefficiens* membrane. In WT, the hopanoid precursor squalene (eluted at ∼14.25 min), diploptene (at ∼16.96 min), diplopterol (at ∼17.97 min), and tetrahymanol (at ∼ 18.85 min) were identified regardless of cumate addition **(Fig. 1E, left panels)**. Pcu-*shc* did not make any detectable hopanoids after EtOH treatment but did produce squalene, whereas cumate restored the production of all types of hopanoids. Curiously, squalene levels were also depleted by cumate addition **(Fig. 1E, right panels)**, likely due to SHC overexpression to levels that fully exhausted the squalene substrate. Together, our assays demonstrate that cumate depletion in the Pcu-*shc* results in complete hopanoid loss and growth inhibition, and that 25µM cumate is sufficient to restore WT lipid composition and growth.

### Hopanoids affect stress tolerance in free-living cultures

Previous work in *B. diazoefficiens* demonstrated that specific subclasses hopanoids, namely the 2-methyl hopanoids and extended hopanoids, are required for survival of membrane-targeting stresses in culture (3, 4, 27); however, whether stress survival is further inhibited by total hopanoid loss has not been investigated in this organism. To address this question, we performed growth curves of Pcu-*shc* and WT strains under three abiotic stresses - low pH, high temperature, and high osmotic pressure - commonly encountered by rhizobia within plant nodules (29). Under low pH conditions (pH = 5.0 compared to the optimal pH=6.6), WT growth was unaffected irrespective of cumate addition (**Fig. 2A**), whereas Pcu-*shc* failed to grow without cumate supplementation. Similarly, Pcu-*shc* cannot survive without cumate under high temperatures (37°C compared to 30°C) **(Fig. 2B)**, but both low pH and high temperature growth could be partially restored by cumate in this strain. Under osmotic stress caused by 5% sucrose, Pcu-*shc* still grew without cumate, albeit slowly, and in this case growth inhibition was fully relieved by cumate (**Fig. 2C**). Dual temperature and osmotic stress roughly phenocopied temperature stress alone, with a slight growth elevation (**Fig. 2D**).

**Figure 2.**
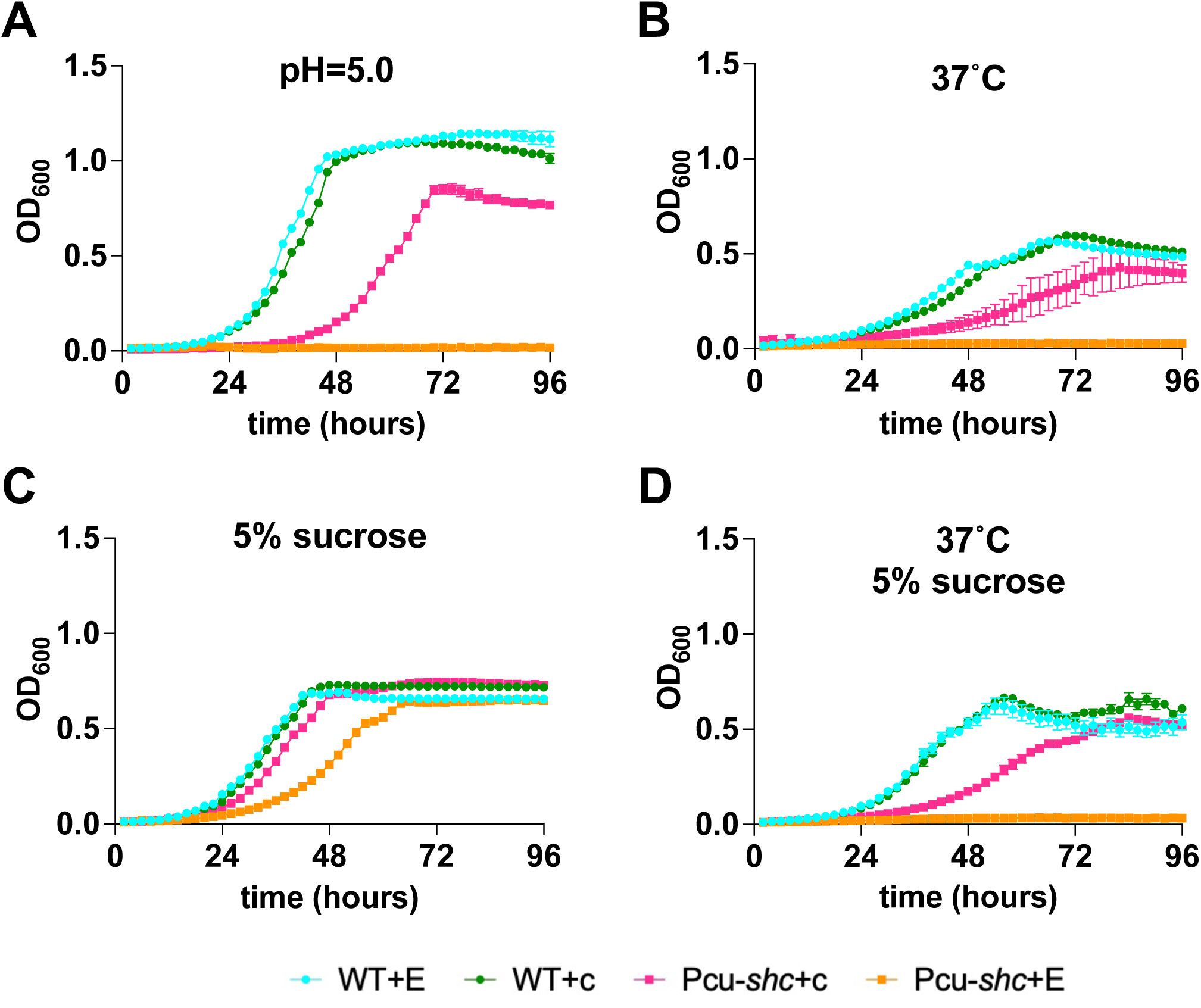
Cumate treatment rescues general stress sensitivity of Pcu-*shc*. 4 day of growth curves of WT and Pcu-*shc* with and without cumate in AG media under **(A)** low pH stress of 5.0, **(B)** high temperature stress at 37°C, **(C)** high osmotic stress of 5% sucrose, **(D)** dual stresses from high temperature and high osmolality.

### Hopanoids participate in soybean-*B.diazoefficiens* symbiosis

Given that hopanoids are essential for *B. diazoefficiens* survival of nodule-relevant stresses in culture, we hypothesized that they may participate in the organism’s native symbiosis with soybean. To test this hypothesis, we inoculated WT or Pcu-*shc* onto soybeans grown in carbon- and nitrogen-free hydroponic growth medium, supplemented with either 25 µM cumate or EtOH. At 27 days post inoculation (dpi), soybeans inoculated with Pcu-*shc* displayed similar nutrient starvation symptoms as non-inoculated (NI) control plants, including uniform foliage chlorosis and significantly stunted growth (**Fig. 3A-B**). Additionally, Pcu*-shc*-inoculated soybeans produced a strikingly lower number of nodules (∼76% less) with reduced nodule dry mass (∼72% lower) compared to plants inoculated with WT **(Fig. 3C-E**). As expected from the low nodule biomass, rates of nitrogen fixation per plant, quantified by the Acetylene Reduction Assay (ARA), were reduced a commensurate amount (∼65%) in the Pcu-*shc-*inoculated soybeans compared to WT-inoculated plants (**Fig. 3F**). However, normalizations of acetylene reduction by nodule number and dry mass were not significantly different between WT- and Pcu-*shc*-inoculated strains (**Fig. 3G-H**), indicating that the reduction of nitrogen fixation in Pcu-*shc* inoculated soybean was due to the decreased nodule number, rather than a core nitrogen fixation defect. An independent biological replicate of these assays was performed that yielded consistent results (**Fig. S2A-H**).

**Figure 3.**
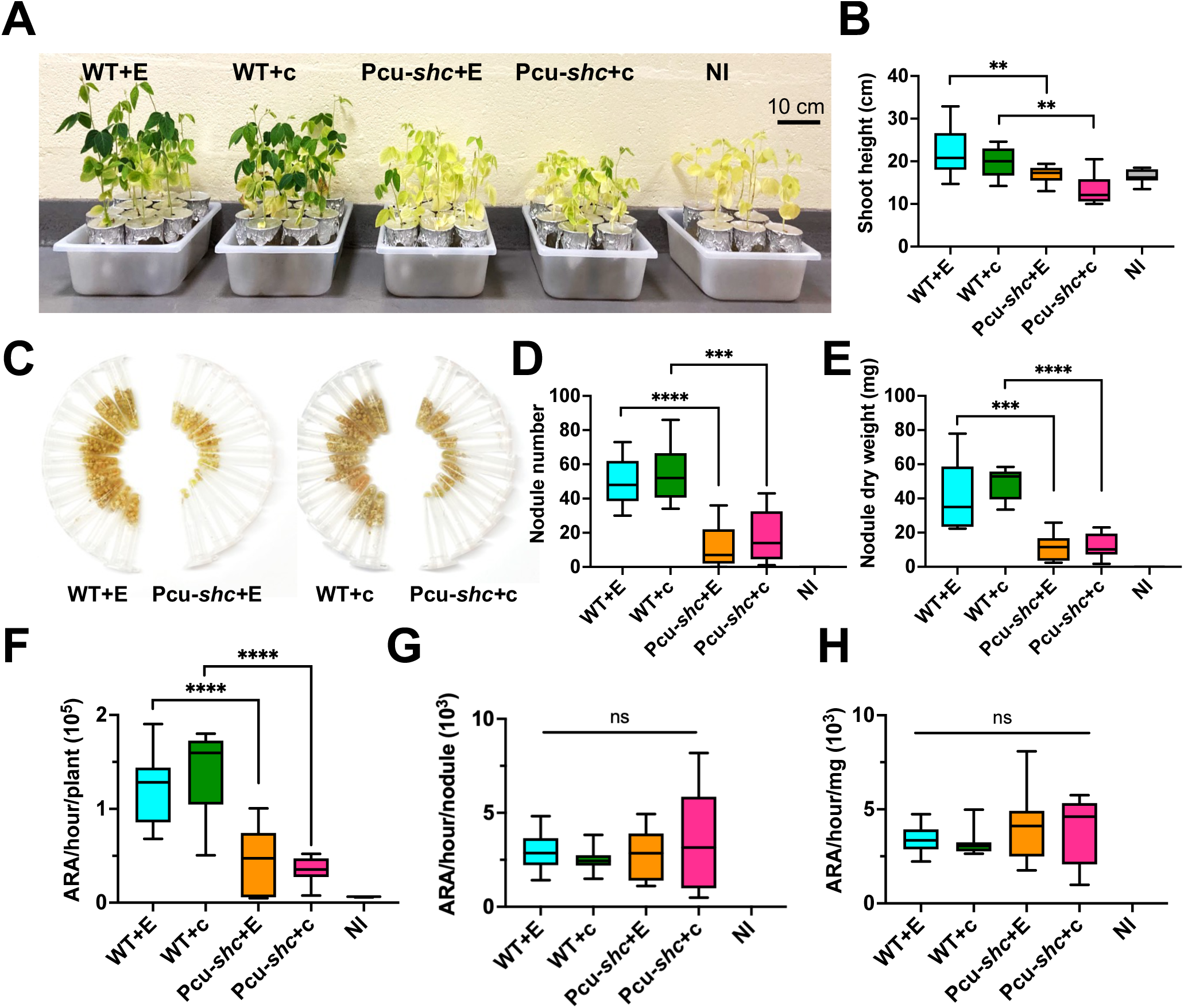
Pcu-*shc* is an inefficient soybean symbiont at 27 days post-inoculation (dpi). **(A)** Comparison of growth soybeans inoculated with WT and Pcu-*shc* with and without cumate respectively (n=9 plants/treatment). Non-inoculated (NI) soybeans (n=7 plants) were grown as control plants. **(B)** Median and quantiles of shoot height (cm) per plant in (A). **(C)** Images of nodules in 1.5 mL Eppendorf tubes collected from soybean in (A). Each tube contains all nodules from a single plant. (**D-E**) Median and quantiles of nodule number and nodule weight (mg) per plant for the treatments of (A). **(F-H)** GC-MS quantification of nitrogenase activity per plant by Acetylene Reduction Assay (ARA). The area of the ethylene peak from each acetylene-treated plant, which is positively correlated with nitrogen fixation, was recorded and normalized by reaction time **(F)**, nodule number **(G)**, and nodule weight **(H).** The median and the quantiles were calculated using GraphPad Prism 9.0. Levels of significance were evaluated using t-test and annotated as follows: P>0.05, non-significant (ns); P<0.01, two asterisks (**); P-value<0.001, three asterisks (***); four asterisks (****). P-value<0.0001.

Interestingly, none of the Pcu*-shc* phenotypes could be restored to WT in plants supplemented with 25 µM cumate (**Fig. 3**). We suspect that cumate was not absorbed into nodule tissues or was metabolized by soybean hosts, though it is also possible that loss of squalene in the Pcu*-shc* plus cumate condition plays a role (**Fig. 1E**). We also examined soybean symbiosis phenotypes at 45 dpi, to determine whether the hopanoid defect could be compensated over time, as shown for extended hopanoid mutants in *Aeschynomene* hosts (4). In line with the results at 27 dpi, soybeans inoculated with Pcu-*shc* grew much slower with shorter shoots, lower nitrogen fixation, fewer nodules, and lower nodule dry weight than WT-inoculated plants (**Fig. S3**), demonstrating that hopanoid phenotypes could not be compensated over this time period.

### Hopanoid mutant-infected soybean nodules are disorganized with low symbiont loads

To assess nodule and bacterial morphologies in symbiosis, we prepared semi-thin (100 µm) sections of ∼20 nodules pooled from 5-10 plants per treatment. Under stereo microscopy, most Pcu*-shc*-infected nodules (95%) had dark pigments indicative of starch granules surrounding the infection zone (**Fig. 4E-H**). Interestingly, some Pcu-*shc* nodules (∼18%) displayed segmented or “patchy” infection zones with starch granules located at the edge of patches (**Fig. 4E, F, H**). In contrast, all the WT-infected nodules had an evenly distributed infection zone pattern across the nodule (**Fig. 4A, C**), even at lower infected cell density (**Fig. 4B, D**), and the frequency of nodules with starch granules was low (9%). These starch granules are not strictly a sign of poor nitrogen fixation; though nearly every Pcu-*shc* infected nodule displayed this phenotype, our previous assays showed that acetylene reduction per nodule dry mass between WT and Pcu-*shc* was not significantly different (**Fig. 3H**). Instead, we interpret it as an indication of an imbalance between photosynthesis of plants and the ability of bacteroids to metabolize them.

**Figure 4.**
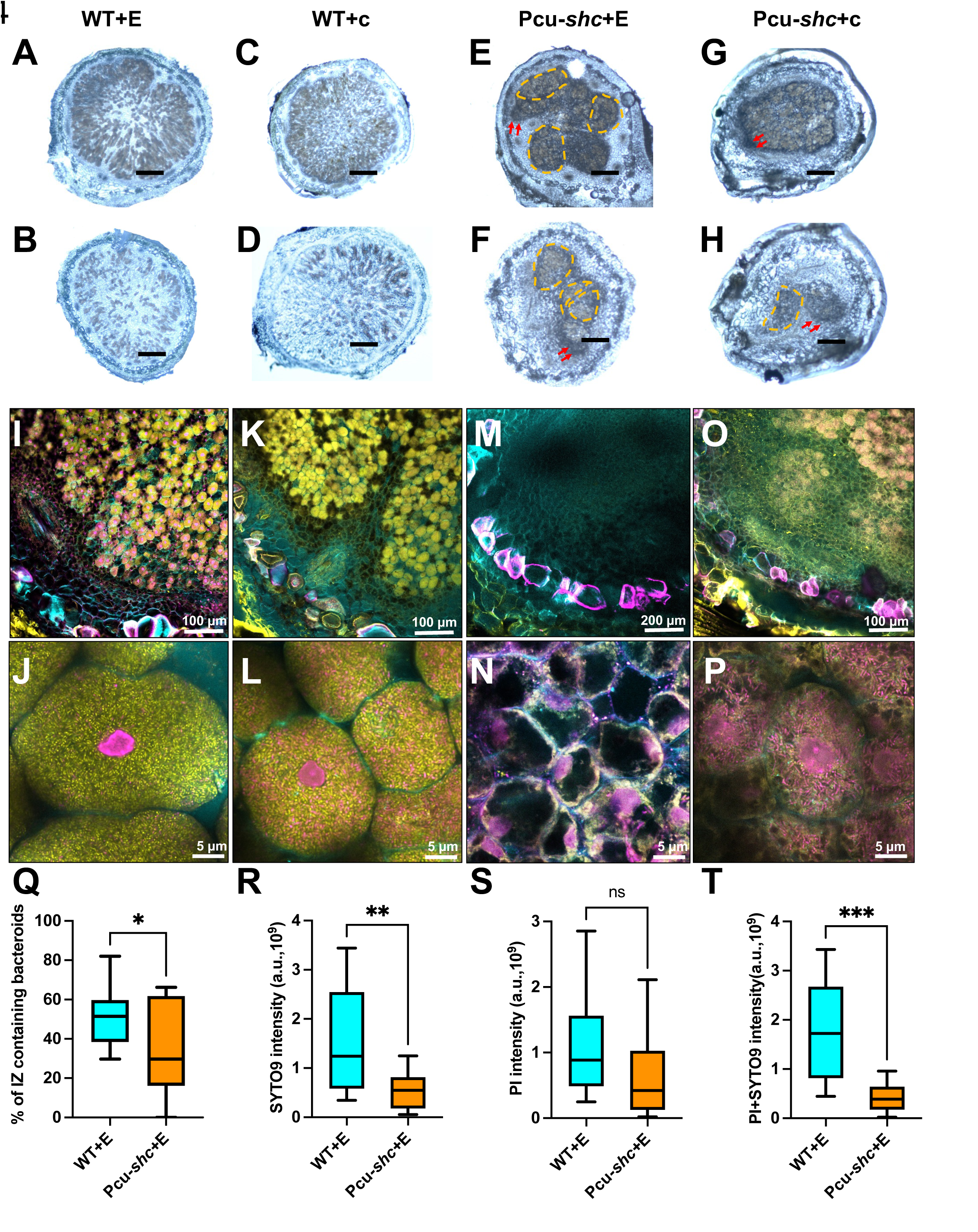
Pcu-*shc* forms some empty nodules on soybean at 27-day post-inoculation (dpi). **(A-H)** Brightfield images of nodule cross-sections obtained by a Zeiss Stemi 508 stereo microscope, scale bars on each image represent 1 mm. Representative images of nodules infected by treatments of WT+E (A-B) and WT+c (C-D), Pcu-*shc*+E (E-F), Pcu-*shc*+c (G-H). The shapes with orange dashed lines are the “patchy” infection zones. The red arrows point to starch granules. **(I-P)** Representative confocal images of nodule cross-sections infected by WT+E (I-J), WT+c (K-L), Pcu-*shc*+E (M-N), Pcu-*shc*+c (O-P), illustrating plant cell walls (Calcofluor, cyan), LIVE bacteria (SYTO 9, yellow), and membrane-compromised bacteria and plant nuclei (propidium iodide-PI, magenta). **(Q-T)** Fiji (ImageJ) quantification of portion of infection zone (IZ) containing “LIVE” (SYTO 9) and “DEAD” (PI) bacteroids (Q) and intensity of LIVE (R), DEAD (S), and total bacterial cells (T), a.u. = arbitrary units. Levels of significance are annotated as in Fig.3

To examine if the loss of hopanoids affected bacteroid viability and persistence, we stained a similar number of nodule cross-sections as above with propidium iodide (a membrane-impermeable DNA dye), SYTO 9 (a membrane-permeable DNA dye) and Calcofluor white (a plant cell wall stain). High resolution confocal imaging with a Zeiss Airyscan revealed all of the WT-infected nodule cells contained a high density of symbionts predominantly stained with SYTO 9 (**Fig. 4I-L, Fig.S4).** In contrast, a portion of Pcu-*shc*-infected nodules (∼30%) contained no or very few symbionts that were predominantly PI-stained **(Fig. 4M-P; Fig.S5)**. We also used Fiji (ImageJ) to quantify the bacteroids occupancy and intensity. It revealed that a larger portion (∼50%) of infection zones (IZ) in WT-infected nodules contained SYTO 9- or PI-stained bacterial cells compared to Pcu-*shc*-infected nodules (∼33%) (**Fig. 4Q)**. Those WT-infected nodules also had stronger intensity of SYTO 9 stained (LIVE) cells and total cells in spite of no-difference PI-stained (DEAD) cells (**Fig.4R-T**). The independent biological replicate of these assays yielded consistent results (**Fig. S2I-L**). Together, these results reflected that Pcu-*shc*-infected nodules had fewer overall bacterial cells with low viability.

In largely empty Pcu-*shc*-inoculated nodules, plasma membranes of soybean nodule cells were separated from their cell walls, suggesting cell death (**Fig. 4N, Fig. S5**). Considering that ARA per nodule and per nodule dry mass were equivalent between WT and Pcu-*shc* infected nodules, we assumed that the low nitrogen fixation of empty or under-occupied Pcu-*shc* nodules were counteracted by WT-like nodules with high occupancy **(Fig. S6)**. This heterogeneity of hopanoids mutant developmental phenotypes may also explain the high variation of ARA per nodule and per mass (**Fig. 3G-H**).

### Total hopanoid loss prevents *B. diazoefficiens* symbiosis with *A. afraspera*

We also examined the symbiotic phenotypes of Pcu-*shc* without cumate on the tropical legume host *A. afraspera*, in which extended hopanoid and 2-methylated hopanoid mutant phenotypes were previously examined (4). In this host, the plant shoot height, the number and mass of nodules infected by Pcu-*shc* was substantially reduced at 28 dpi (**Fig.S7A-D**), and all of these mutant-infected nodules were much smaller than WT (**Fig.S7E-F**). Although only a few nodules were present per plant, nodule sections stained as above suggest all nodules were empty or contained a low density of undifferentiated bacteria (**Fig.S7G**).

### Transcriptomic studies suggest that hopanoids suppress the expression of flagellar motility-related genes and enhance expression of genes involved in protein secretion

To understand the molecular mechanisms underlying the phenotypes of Pcu-*shc* under hopanoid-free conditions, whole-genome transcriptomics of free-living WT and Pcu-*shc* with and without cumate were investigated. Normalized read counts of WT and Pcu-*shc* per treatment from two biological replicates with three technical replicates each (with the exception of WT with EtOH only, which has only 5 replicates total, as one sample contained poor quality RNA) are provided in **Table S2.** We obtained 25-50 million reads per sample with 200x-400x coverage, and 96% of the reads could be aligned to the *B. diazoefficiens* USDA110 genome. In all samples a large majority of the genes (98%) had non-zero counts.

As expected from our qRT-PCR assays **(Fig. 1C)**, the coverage and average of read counts of *shc* in Pcu-*shc*+E (160±6) was ∼65-fold lower than WT+E (10580±460), whereas they were ∼7-fold higher in Pcu-*shc*+C (80841±3043) compared to WT+C (10064±345) **(Fig. 5A, Table.S2)**. Also, as expected from our growth curves and symbiosis phenotypes, the Pearson correlation coefficient between Pcu-*shc*+E replicates with all other samples was lower (∼0.94, in blue) than the coefficients of correlation between all other samples (∼0.99 in orange) **(Fig. S5)**.

**Figure 5.**
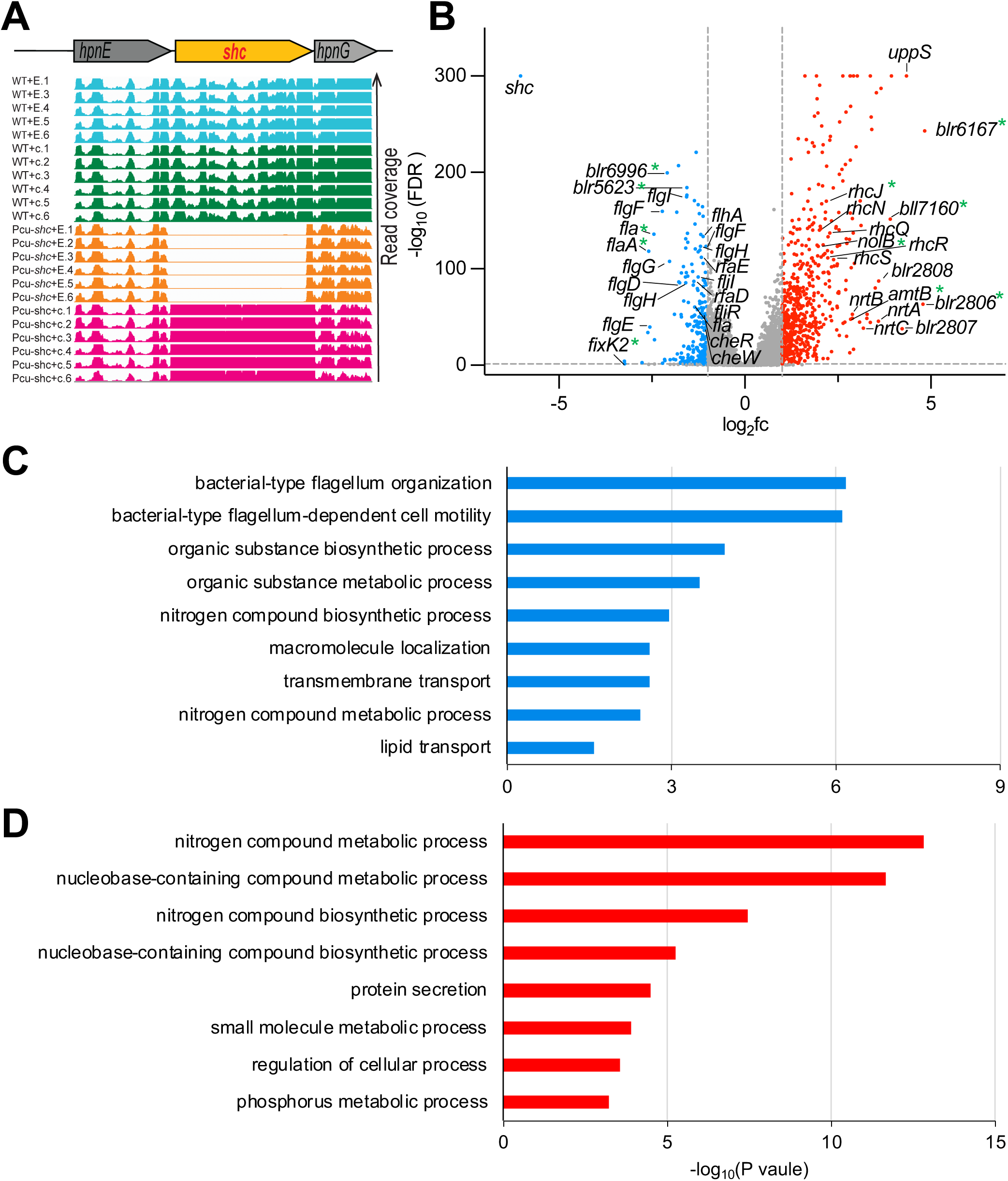
Pcu-*shc* transcriptomics suggests defects in motility and chemotaxis and enhanced nitrogen metabolites and protein secretion. **(A)** Read coverage of the *shc* genomic region in WT and Pcu-*shc* with and without cumate. **(B)** Volcano plot of all genes based on read fold change (fc) and false discovery rate (FDR) of Pcu-*shc*+E compared to WT+E. The red dots represent up-regulated genes (log_2_fc≥1, FDR <0.05), the blue dots represent down-regulated genes (log_2_fc≤-1, FDR<0.05), and the gray dots represent non-differentially expressed genes (−1<log_2_fc<1, FDR≥0.05). The genes with asterisks are confirmed by qRT-PCR in Table.S6 **(C-D)** Go analysis of down-regulated (C) and up-regulated (D) in biological process categories.

We performed differential gene expression (DEG) analysis on transcripts from Pcu-*shc*+E compared to WT+E **(Table S3)**, where DEGs in Pcu-*shc*+E were defined by a log_2_foldchange ≥1 or ≤-1 and a false discovery rate-adjusted p-value ≤0.05. This analysis identified 595 up-regulated DEGs (**red, Fig. 5B**, **Table S4**) and 247 down-regulated DEGs **(blue, Fig. 5B, Table S5)**, accounting for 7% and 3% of the genome, respectively. The majority of DEGs were uncharacterized genes and a heatmap of the top 20 up- and and down-regulated DEGs is attached in **Fig. S6.** To further assess the quality of the RNA-seq data, 12 genes from the up- and down-regulated DEGs (6 up-regulated genes, 5 down-regulated gene, 1 non-significance gene) were quantified by qRT-PCR. The results showed all the genes were consistent with the RNA-seq data (**Table S6**), confirming the fidelity of our RNA-seq data for further analysis. Among down-regulated DEGs, many genes are involved in the regulation of flagella (*e.g.*, *fla*, *flaA*), flagellar assembly (*e.g.*, *flaF*, *flaE)*, and chemotaxis (*e.g.*, *cheR*, *cheW*), which are highlighted in the volcano plot (**Fig. 5B**). GO-term enrichment analysis also confirmed that flagellum organization and flagellum-dependent motility were the top GO terms associated with down-regulated DEGs (**Fig. 5C**). Genes involved in lipopolysaccharide (LPS) biosynthesis and symbiosis, such as *rafD* and *rafE* (30), also are among the down-regulated DEGs (**Fig. 5B**). This may reflect the role of hopanoids as a structural component of the lipid A moiety of LPS in *Bradyrhizobium* (5).

GO term enrichment analysis indicated up-regulated genes are most strongly associated with nitrogen metabolic processes (**Fig. 5D**). These include the ammonium transporter *amtB* and each gene in the operon *blr2803-2809*, which is involved in NO^3-^ assimilation. Proteins encoded include: an ABC-type NO^3^-transport system NrtABC (blr2803–05), a major facilitator superfamily (MFS)-type NO^3^-/NO^2^− transporter (Blr2806), bacterial hemoglobin (Blr2807), an FAD-dependent NAD(P)H oxidoreductase (Blr2808), and the catalytic subunit of the assimilatory NO^3−^ reductase NasA (Blr2809). The most up-regulated gene in Pcu-*shc*+E, *uppS*, is an isoprenyl transferase involved in terpenoid biosynthesis, upstream of hopanoid biosynthesis. Genes involved in secretion systems, such as *hlyD* and *bll6293* of the type I secretion system (T1SS) and an operon composed of 25 type III secretion system (T3SS) genes (*e.g.*, *rhcJ*, *rhcN*, *rhcQ*, *nolB*, *rhcS*, *rhcR* etc), are the second major category of up-regulated DEGs in Pcu-*shc*+E **(Fig. 5B, 5D**). Recent research revealed that mutants in secretion systems in plant-beneficial bacteria modulate prokaryotic and eukaryotic interactions in the rhizosphere (31)

### Hopanoid depletion prevents swimming motility

To determine whether reduced expression of flagellar motility-genes was sufficient to inhibit Pcu-*shc*+E motility, swimming assays were in semisolid agar plates. The results showed that WT swimming was unaffected by cumate (**Fig. 6A**), with a motility halo at 12 dpi that was 2-fold larger than at 2 dpi. Pcu-*shc*+E inoculum did not appear to swim between 2-12 dpi, whereas the cumate-complemented strain expanded similarly to WT (**Fig. 6B**).

**Figure 6.**
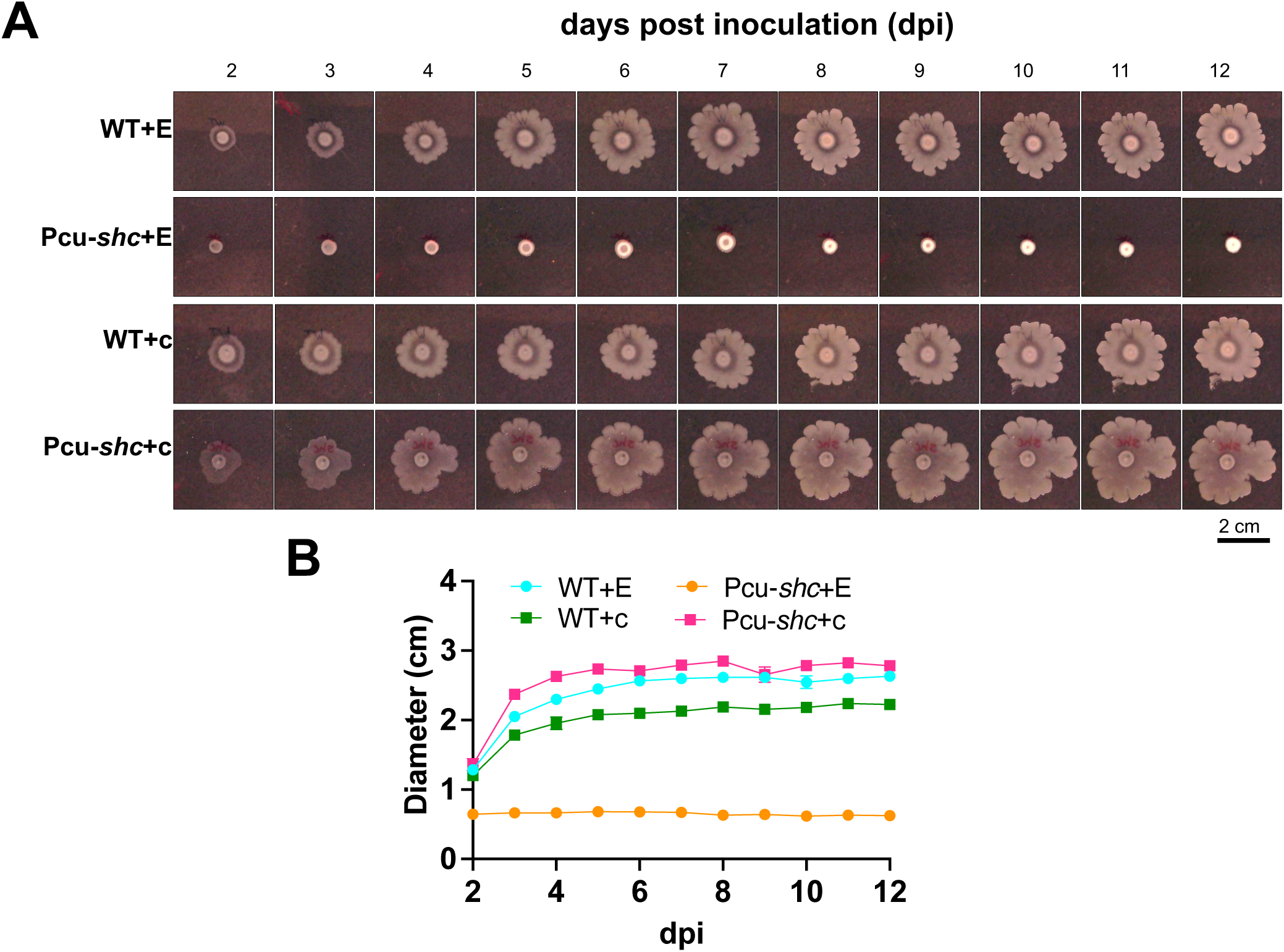
Pcu-*shc* displays compromised flagellar motility. Swimming motility of WT and Pcu-*shc* with and without cumate in starving agar plates (AG without yeast but with 0.1g/L arabinose and gluconate). **(A)** The morphologies of WT and Pcu-*shc* with and without cumate on starvation medium plates from 2-12 dpi. **(B)** The mean ± standard errors (SEM) of diameters of colonies from the four treatments in (A). N = 3 technical replicates for each treatment. The error bars are low that they are covered by the symbols of each data set.

## DISCUSSION

Relatively few studies explore the biological function of bacterial lipids on plant-microbe interactions. A difficulty of such work is the construction and validation of lipid biosynthesis mutants. This validation typically requires lipid extraction and quantification by mass spectrometry, a technically challenging approach that is less widely available than methods for tracking protein depletion. Here we provide a thorough validation of an inducible total hopanoids mutant strain, previously generated by colleagues through a clever genetic approach to bypass the difficulty of selecting for a simple *shc* knockout (24). We demonstrate that the strain is suitable for cumate dose-dependent manipulation of not only *shc* expression but also membrane hopanoid content in a pure culture setting.

### Consequences of hopanoid loss in stress survival

In the absence of hopanoids, *B. diazoefficiens* stress sensitivity is markedly more severe than reported in other hopanoid mutants. Under pH stress at pH 4.5-5.0, Δ*shc* strains of *Rhodopseudomonas palustris* TIE-1 (27) and *Burkholderia cenocepacia* K56-2 (10) have delayed growth but are still viable, whereas the equivalent total hopanoid knockout of *B. diazoefficiens* cannot grow. This may suggest that hopanoids are more central to pH stress resistance in our organism, consistent with previous work on extended hopanoid mutants (Δ*hpnH*) of *B. diazoefficiens* (3, 24). However, we cannot discount the effects of distinct medium compositions used to cultivate the different species.

In the context of temperature stress, the Δ*hpnH* mutant of *B. diazoefficiens* also cannot grow at 37 °C (3) suggesting extended hopanoids are the primary contributors to this phenotype in the Pcu-*shc* strain. This Δ*hpnH* strain has more mild growth inhibition under osmotic stress (24) than Pcu-*shc*+E, and the 2-methylated hopanoid mutant Δ*hpnP* has no osmotic stress phenotype (3), indicating that the short hopanoid class is the most relevant to osmotic stress survival. Overall, the stress sensitivity phenotypes of Pcu-*shc*+E under our culturing conditions are more severe than previously published mutants. The strong Pcu-*shc*+E phenotypes, combined with its inducibility, demonstrate that it is a strong model for future molecular studies of hopanoid-mediated stress resistance.

### Relative effects of hopanoids across *Bradyrhizobium*-legume symbioses

The consequences of hopanoid loss in *Bradyrhizobium*-legume symbioses have now been examined in three models: (i) *B.* BTAi1 Δ*shc* in *A. evenia* (5), (ii) *B. diazoefficiens* Δ*hpnH*, Δ*hpnP*, and Pcu-*shc* in *A. afraspera* (3, 4), and (iii) *B. diazoefficiens* Δ*hpnH*, Δ*hpnP*, and Pcu-*shc* in soybean (3). In cases where hopanoid mutants negatively affected symbiosis, but did not block symbiosis completely, the phenotypes are nearly identical. *B.* BTAi1 Δ*shc* in *A. evenia*, *B. diazoefficiens* Δ*hpnH* in *A. afraspera*, and *B. diazoefficiens* Pcu-*shc* in soybean all had a roughly two-fold lower rate of acetylene reduction per plant than the WT strain at 2-4 weeks post-inoculation, with a subset of nodules exhibiting low symbiont load, disorganization of the central infection zone, and signs of early nodule senescence or nodule cell death (3–5), What differs between the three models is which hopanoids must be depleted to yield this common phenotype. In *A. afraspera* hosts, total hopanoid loss prevents symbiosis entirely, but the extended hopanoid knockout Δ*hpnH* yields the common phenotype, suggesting that short hopanoids are most important in the *B. diazoefficiens*-*A. afraspera* interaction. In soybean and *A. evenia*, the common phenotype is recapitulated in a total hopanoid knockout, though we note that Δ*hpnH* and Δ*hpnP* were not tested in *A. evenia*.

Why total hopanoid loss elicits a stronger phenotype in the *B. diazoefficiens*-*A. afraspera* symbiosis is not clear. It is possible that the types of stresses associated with this host are different from those of *A. evenia* and soybean, but we find this conclusion unlikely. Instead, we believe that the stronger phenotype of *B. diazoefficiens* Pcu-*shc* in *A. afraspera* reflects the fact that these two species do not interact in nature. Though *B. diazoefficiens* can engage in this symbiosis, it is less efficient than the native symbiont *B.* ORS285 (32), suggesting that *B. diazoefficiens* has not been adapted to survive in the *A. afraspera* tissue. Regardless, the similarity of the total hopanoid knockout phenotypes of the native *B.* BTAi1-*A. evenia* and *B. diazoefficiens*-soybean pairings confirm that hopanoids are indeed important for natural *Bradyrhizobium*-legume symbiosis. All studies also agree that 2-methyl hopanoids produced by HpnP are not major contributors to symbiosis, and instead that short and extended hopanoids are the more significant lipids in this context.

### Molecular mechanisms of hopanoid function

Prior analysis of the *B. diazoefficiens*-*A. afraspera* symbiosis proposed that the reduced nitrogen fixation of hopanoid mutants is a product of at least two independent defects: delayed initial infection of plant roots, perhaps relating to reduced strain motility, and slower root nodule growth, perhaps due to lower levels of symbiont proliferation and survival *in planta* (4). This manuscript supports a similar model in soybean. We observed both reduced flagellar assembly gene expression and loss of swimming motility in the Pcu-*shc*+E condition, consistent with the reduced swimming motility of *B. diazoefficiens* Δ*hpnH* (4) and B. *cenocepacia* Δ*shc* (10), as well as lower stress resistance in culture and bacteroid densities in nodules. Our transcriptomics work introduces several new potential mechanisms by which hopanoids depletion inhibits symbiosis: excessive nitrogen assimilation, altered secretion of symbiotically-relevant effector proteins, and loss of chemosensation.

How hopanoids affect these processes, at the molecular level, is not clear. Some of the observed phenotypes may arise from activation of general stress response (GSR) mechanisms in hopanoid mutants. In *R. palustris* TIE-1, expression of 2-methyl hopanoids is regulated by the extracellular function (ECF) sigma factor, which mediates GSR in this organism (33). Though it has not been tested directly, it is possible that the levels of some or all hopanoids is also linked to GSR activation in *B. diazoefficiens*, or to the activation of membrane-specific stress response pathways. Another hypothesis is that hopanoids affect membrane-based processes through their role in biophysically reinforcing or compartmentalizing cell membranes reviewed in (34). Bacterial flagellar motility, chemotaxis, protein secretion, and the nitrate assimilatory pathway all involve large membrane-associated protein complexes that may rely on proper membrane mechanics to function properly. There is some evidence for this hypothesis in the case of protein secretion: the Sec translocon ATPases of *E. coli* (35) and *Streptococcus pneumoniae* (36) are sensitive to membrane lipid composition and biophysics. This possibility will be an exciting topic of future investigation.

### Challenges for future tool development

As we’ve demonstrated, the inducible mutant strategy used to develop the Pcu-*shc* strain (24) is an extremely useful workaround for generating mutants of *B. diazoefficiens* genes that are essential or otherwise difficult to delete cleanly. However, it has limited use *in planta*, as we could not rescue *shc* depletion phenotypes in the host. Further tool development to identify tissue-permeable inducer molecules, and the identification of promoters that rescue *Bradyrhizobium* gene expression at specific stages of nodule development as have been done for *Sinorhizobium meliloti* (37) would significantly enhance our ability to understand bacterial gene function in *Bradyrhizobium*-legume symbioses.

## METHODS

### Bacterial strains and cultivation

Wild-type *Bradyrhizobium diazoefficiens* strain USDA110 spc4 (WT *B.diazoefficiens* USDA110) originally was obtained as a gift from Dr. Hans-Martin Fischer (ETH Zurich) to Dr. Dianne Newman (Caltech), who then gifted the strain to Dr. Belin. Hopanoid mutant strain Pcu-*shc*::*Δshc* was generated as described (24) and provided as a gift from Dr. Dianne Newman (Caltech).

For all experiments, cells from 10% glycerol stocks stored at −80°C were streaked on agar plates made with rich AG medium (4.6 mM sodium gluconate, 6.6 mM arabinose, 1 g/L yeast extract, 6 mM NH4Cl, 5.6 mM MES, 5 mM HEPES, 1 mM Na_2_HPO_4_, 1.76 mM Na_2_SO_4_, 88 µM CaCl_2_, 25 µM FeCl_3_, 0.73 mM MgSO_4_, pH 6.6). Plates were grown aerobically at 30°C until single colonies appeared, typically for 3-5 days, and colonies were selected for inoculation into 5 mL liquid rich AG. Liquid cultures were grown to OD_600_ = 0.5-0.8 under aerobic conditions in Eppendorf Innova incubating shakers at 30°C and 250 rpm. Culture tube OD_600_ values were measured using a Thermo Scientific GENESYS 30 Visible Spectrophotometer with a test tube adapter module.

For cultivation of Pcu-*shc*::*Δshc*, growth plates were supplemented with 25 µM cumate (4-Isopropylbenzoic acid; Millipore Sigma SKU 268402) diluted from a 25 mM (1000X) stock solution in EtOH. Initial cultures of Pcu-*shc*::*Δshc* and WT were grown in rich AG without cuamte to OD_600_ = 0.5-0.8. These cultures were subcultured to OD_600_ = 0.02 and grown to OD_600_ = 0.5-0.8, in either 0.1% EtOH only (for minus cumate samples) or 0.1% EtOH with 25 µM cumate (for plus cumate samples). Both 25 mM (1000X) cumate/EtOH and EtOH stock solutions were sterilized by passing through 0.22 µm filters and stored at −20°C.

### qRT-PCR

qRT-PCR was performed as previously described with minor modifications (38). Briefly, total RNA from WT or Pcu-*shc*::Δ*shc* strains at early exponential phase (OD_600_ = 0.5-0.8) was extracted using a RNeasy® Mini Kit (Qiagen) according to the manufacturer’s recommended protocol. The genomic DNA was removed using on-column DNase-I digestion (Qiagen) for 15 min. RNA concentration and purity were determined using a Nanodrop^TM^ one (Thermo Scientific). Total RNA (500 ng) was reverse transcribed using random primers (Promega) and AMV Reverse Transcriptase (Promega) according to the manufacturer’s protocol.

Quantitative PCR (qPCR) with Fast SYBR Green® PCR Master Mix (Thermo Fisher) was performed using the CFX96^TM^ Real-Time System (Bio-Rad) in Optical 96-Well Fast Plate (Stellar Scientific). Each reaction contained 5 ng cDNA and 250 nM gene-specific primers. The relative expression of the target genes was normalized to the 16S rRNA housekeeping gene based on the mean of cycle threshold (Ct) values and ΔΔCt method. All values are the means ± standard errors of three replicates. The primers for qRT-PCR are listed in **Table. S1**.

### Growth curves

Three individual colonies from WT and Pcu-*shc*::Δ*shc* strains on agar plates were picked up and cultivated in 5 mL AG media, as noted above. Once the bacterial strains reached early exponential phase (OD_600_=0.5-0.8), the culture OD_600_ values were adjusted to the same value in fresh AG medium. Cultures were then diluted 1:100 into 200 µl fresh AG medium in individual wells of a 96-well plate. For growth curves of WT and Pcu-*shc*::Δ*shc* under variable cumate concentrations, AG media was supplemented with either 0.1% EtOH only (for minus cumate samples) or 0.1% EtOH with 5,10, 25, 50, or 100 µM of cumate (for plus cumate samples). For growth of the strains under stress, AG medium was supplemented with 5% sucrose or adjusted to pH 5.0. Each well was overlaid gently with 30 µl of mineral oil to prevent evaporation.

The plate was placed into a microplate reader BioTek EPOCH 2 set to 30°C (most conditions) or 37°C (heat stress conditions) with continuous shaking. The OD_600_ was measured every two hours for 96 hours. Data were plotted in Prism GraphPad.

### Hopanoid extraction and quantification by GC-MS

Hopanoid lipids were purified and quantified as previously described with minor modifications (27, 39). Briefly, 5 mL cultures of WT and Pcu-shc::Δ*shc* were harvested in early exponential phase by spinning down in a swinging bucket centrifuge at 3,250 x g for 30 min. Cell pellets were resuspended in 50 uL of water and transferred into 9 mm Clear Glass Screw Thread Vials (1.7 mL Tapered Base,Thermo Scientific™ SKU C40009) with Vial Caps (Thermo Scientific™, SKU 60180-516). MeOH (125 uL) and CH_2_Cl_2_ (62.5 uL) were added in sequence in a 2:1 ratio, and the vials were capped and sonicated in a Branson M2800H Ultrasonic Bath (40 kHz, Power 110 W) for 30 min at room temperature. The total lipids were extracted by addition of ∼200 uL CH_2_Cl_2_ and mixed by pipetting. After the sample settled for 2-4h at room temperature, permitting the organic and aqueous layer to separate, the lower organic layer that contained the hopanoids was collected, dried in a 60 °C oven, and acetylated in 100 uL of 1:1 pyridine: acetic anhydride (Ac2O) for 30 min at 60 °C.

Acetylated hopanoids were analyzed on a Thermo Scientific TRACE 1300 gas chromatograph coupled with an ISQ 7000 single quadrupole mass spectrometer. The protocols were modified from those previously described (27, 40). One microliter of acetylated hopanoids was injected into the gas chromatograph and run through a Restek^TM^ Rxi^TM^-XLB column (30 m x 0.25 mm x 0.10 um, Restek 13708) with an injector temperature of 325°C and in split mode (4:1) with helium as the carrier at constant flow of 1.5 ml/min. The GC oven was programmed as follows: 100°C retention for 2 min then ramped at 15°C/min to 320°C with a 5 min hold. The mass spectrometer was operated in full scan mode over 50–750 amu at 70 eV in electron impact (EI) ionization mode. The MS transfer line temperature was held at 325°C and the ion source temperature at 250°C. Hop-22(29)-ene (Sigma-Aldrich, PubChem Substance ID: 32974796), also known as the short hopanoid diploptene, was used as standard.

Data collected from GC-MS were processed by the Chromeleon™ 7.3 Chromatography Data System (CDS) Software. The compounds were identified by comparison with the standard in terms of retention times, NIST compound library hits identified in Chromeleon, and mass spectra molecular ions, as described previously (3, 5, 27, 40–42).

### Soybean and *Aeschynomene* cultivation and inoculation

*Aeschynomene afraspera* (*A.afraspera*) seeds were produced in-house and cultivated as previously described (4). Soybean (*Glycine max*) seeds (non-GMO forage variety) were purchased from the Hancock Seed Company. Seeds were sterilized by rinsing in 95% EtOH, incubating in 5% bleach for 5 minutes on a nutating shaker, and washing 5 times in ultrapure water. Using sterile metal forceps, cleaned with ethanol periodically in a biosafety cabinet, sterilized seeds were placed on freshly poured 1% water/agar plates. The plates were sealed with parafilm and covered in aluminum foil to protect from light. Seeds were germinated in the dark at 30°C for 72 hours.

For planting, 7-9 seedlings for each treatment (WT+EtOH, WT+cumate, Pcu-*shc*+EtOH, and Pcu-*shc*+cumate, non-inoculated control) were guided into sterile 400 mL borosilicate glass beakers (soybean assays) or 250 mm x 25 mm /100 mL large tubes (*A.afraspera assays*) through perforated aluminum foil caps. Beakers or tubes were filled with autoclaved buffered nodulation medium (BNM) modified from the previous study (43) with adjustment to pH 6.5 with ∼ 1 mL of 10M KOH per liter. The BNM medium contained 2 mM Cacl-2H_2_0, 2 mM MES buffer, Nod major salts (0.5 mM KH_2_PO_4_, and 0.5 mM MgSO_4_), Nod minor salts I (16 µM ZnSO_4_*7H_2_0, 50µM H_3_B0_3_, 50µM Mncl_2_*4H_2_0), Nod minor salts II (1µM Na_2_MoO_4_*2H_2_0, 0.1µM CuS0_4_*5H_2_0, 0.1 µM CoCI2*6H2O), Fe-EDTA (50 µM Na2EDTA, 50µM FeSO_4_*7H_2_0). An additional 500 µM KNO_3_ was added to provide the minimum nitrogen source required for soybean growth before nodule formation. Seedlings were grown for 5-7 days in Conviron GEN2000 chambers with light intensities of 600 µmol/m^2^/s at 28°C and 80% RH for a 16:8 hour day:night cycle.

WT and Pcu-*shc*::*Δshc* cultures were grown to OD_600_ = 0.5-0.8 in AG media without cumate and then subcultured with or without cumate, as described above, to OD_600_ = 0.5-0.8. Bacterial cultures were pelleted at 3520 x g for 30 minutes at room temperature and resuspended in rich AG to a final OD_600_ = 1.0. After adjusting the culture OD_600_, 4 mL or 1 ml of culture suspension was added to each soybean or *A.afraspera* respectively, along with 400 µL of 25 mM (1000X) cumate, or 400 µL of EtOH only. Plants inoculated with different strain/cumate combinations were cultivated in separate bins to prevent cross-contamination. After inoculation, plants were maintained in the GEN2000 chambers until harvesting for nodule analysis at 27-28 days of post inoculation (dpi) or 40-45 dpi under the same conditions described above. In longer experiments (exceeding roughly 30 days post-inoculation), plants were replenished with BNM as required.

### Acetylene reduction assays and plant phenotyping

At the time points indicated in the results, soybeans were cut at the hypocotyl with sterile razor blades, and the aboveground tissues were used for measuring the shoot height while the roots were placed inside 250 mL Balch-type serum vials containing ∼10 mL of ultrapure water. For *A.afraspera* assays, the whole plants were put into vials and shoot heights were measured after acetylene reduction assay. Vials were sealed using bromobutyl rubber stoppers and aluminum crimp caps, and 10 mL of headspace was removed and replaced with 10 mL acetylene gas (Airgas USA) using a sterile syringe with a 16G hypodermic needle. The ARA reaction start time was recorded immediately after acetylene injection. Vials were then inverted to verify a gastight seal was maintained, based on the absence of visible gas bubbles in the water. Vials containing air only or non-inoculated plants were also treated with acetylene as controls.

Acetylene-treated plants were maintained overnight in GEN2000 chambers and serially removed for headspace analysis by GC-MS. Gastight syringes with removable 22s G, 2 inches, point style #2 needles (Hamilton Company) were used to inject 100 µL headspace samples into TRACE 1300/ISQ 7000 GC-MS equipped with a TracePLOT TG-BOND Q+ column (30 m x 0.32 mm x 10 µm, Thermo Scientific, Cat. No. 26005-6030) with an injector temperature of 100°C in splitless mode, with helium as the carrier at constant flow of 1.5 ml/min. The GC oven was programmed with constant 60°C. The mass spectrometer was operated in 20–100 amu at 70 eV in electron impact (EI) ionization mode. The MS transfer line temperature was held at 250 °C and the ion source temperature at 250 °C.

The compound peak areas and ARA reaction stop time were recorded. Acetylene and ethylene were identified by searching the NIST compound library in Chromeleon. The ARA reaction duration was hand calculated. The areas of the ethylene peak were calculated and the acetylene reduction rates were calculated as:

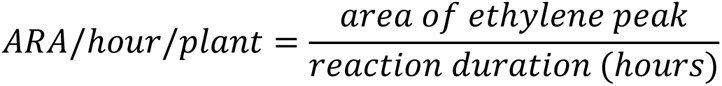

After measuring acetylene reduction, plants were removed from the bottles and the nodules were counted and carefully harvested using a razor blade. The harvested nodules were transferred into pre-weighed Eppendorf tubes and were photographed, and then dried at a temperature of 60°C for at least 48 hours before weighing to calculate their dry mass. For reporting acetylene reduction per nodule and per milligram of nodule dry weight, the formulas below were used:

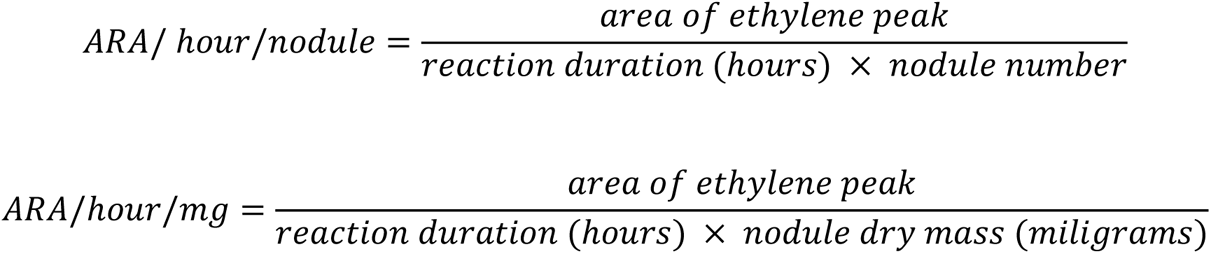

### Nodule staining and microscopy

Root nodule semi-thin sections (100 µm thickness) were collected using a 7000smz-2 vibratome (Campden Instruments). Brightfield images of nodule sections were taken using a Zeiss Stemi 508 stereo microscope with an Axiocam 305 color camera. For fluorescence images, nodule sections were immediately stained with 5µM SYTO 9 and 30µM propidium iodide (PI) using a LIVE/DEAD *Bac*Light Bacterial Viability Kit (Thermo Fisher Scientific Cat. No. L13152) according to the manufacturer’s protocol. After LIVE/DEAD staining, cells were fixed in PBS containing 3.7% paraformaldehyde (Sigma-Aldrich, SKU F8775) for 30 minutes at room temperature. Fixed sections were washed 3X in PBS and transferred to a solution of PBS with 0.01% Calcofluor stain (Fluorescence brightener 28 disodium salt solution; Sigma-Aldrich, SKU 910090). Samples were stained at room temperature for 30 minutes or overnight at 4°C and then washed 3X in PBS. Samples were then mounted onto slides in one-well, 20 mm diameter, 0.12 mm deep Secure-Seal spacers (Thermo Fisher Scientific Cat. No. S24736) in Fluoromount-G^TM^ mounting medium (Thermo Fisher Scientific, Cat. No. E141473) and were sealed with 20 mm square #1.5 coverslips and Seche Vite fast dry nail polish (Amazon). The slides were cured at 4°C overnight prior to imaging.

High resolution images were taken using a ZEISS LSM 980 confocal microscope with an Airyscan 2 super-resolution module using 10X (0.45 NA) or 63X (1.4 NA) objectives. The images were acquired by brightfield or in fluorescence mode with the following filters for each dye: Calcofluor - 405 nm laser excitation and 422-477 nm emission; SYTO 9 - 488 nm laser excitation and 495-550 nm emission (“LIVE” stain); propidium iodide (PI) - 561 nm laser excitation with 607-735 nm emission (“DEAD” stain). All fluorescence images were acquired using GaAsP-PMT detectors. Quantitative analysis of nodule images was performed using custom ImageJ macros.

### RNA-seq library construction and analysis

Three individual colonies of WT and Pcu-*shc*::Δ*shc* growth with and without 25 µM cumate were cultured to early exponential phase and total RNAs were extracted by Qiagen RNeasy Mini Kit as described previously (44). RNA quantification was achieved using a Nanodrop one (Thermo Scientific) and RNA quality was monitored with an Agilent 2100 Bioanalyzer using an RNA 6000 Pico Kit (Agilent Technologies). Ribosomal RNA (rRNA) was depleted and cDNA libraries were prepared from 100 ng total RNA by Stranded Total RNA Prep Ligation with Ribo-Zero Plus (Illumina). The quantity and quality of the prepared library were analyzed by the Qubit dsDNA BR Assay Kit and the Agilent 2100 Bioanalyzer using a DNA 1000 Kit, respectively. A library of samples (2 ng/sample) under each condition (30Mb read/each sample) was loaded on the Illumina Nextseq 500 system with a NextSeq 500/550 High Output Reagent CartriDEG v2 (Illumina) for single-end reads at Carnegie Embryology Department’s in-house sequencing facility. Two independently biological replicates were performed.

Raw data from the two independent experiments were analyzed together by the nf-core/rnaseq pipeline built using Nextflow (45). Based on gene annotation of *Bradyrhizobium diazoefficiens* USDA 110 in EnsemblBacteria (https://bacteria.ensembl.org/index.html), reads were mapped to the genome resulting in a compressed binary version of the Sequence Alignment Map (BAM files) and WIG files for reads visualization. Differential gene expression (DEG) analysis of Pcu-*shc*::Δ*shc* compared to WT was performed using DESeq2 in RStudio (46, 47). To make accurate comparisons of gene expression between samples, raw counts were normalized by the DESeq2’s size factor and batch factor of each sample to correct for variability of the sequencing depth and batch effect from independent experiments. The ratios of normalized counts of each gene in Pcu-*shc*::Δ*shc* and WT were computed as a fold change (fc). Genes with differences in expression (false discovery rate (FDR) < 0.05; log_2_fc was ≥ 1 or ≤ −1) were further used to construct the volcano plots by GraphPad Prism 9.0. The Pierson correlation and heatmap of top 20 up and down-regulated DEGs were constructed by ImageGP (48) (https://www.bic.ac.cn/ImageGP/). The complete dataset from this study has been deposited in the National Center for Biotechnology Information Gene Expression Omnibus with the Accession no.XXXX (note: the accession no. will be available after the acceptance of this manuscript). The Gene ontology (GO) functional enrichment analysis of the up-regulated gene sets and down-regulated gene sets was performed using PANTHER (49, 50)(http://pantherdb.org/).

### Swimming motility assays

Swimming motility assays were performed as previously described with some modifications (51). Three single colonies of each bacterial strain were grown to turbidity (OD_600_=1.15-1.3) in 5 ml of AG media, then diluted to an OD_600_ of 0.02 in 10 ml of fresh AG containing 0.5 g/L of arabinose and gluconate without yeast extract. Cultures were grown to the early exponential phase (OD_600_ = 0.25-0.4) and then diluted to an OD_600_ of 0.05 in fresh starvation media (AG without yeast extract but with 0.1g/L arabinose and gluconate). For each bacterial strain, 5 μl of the adjusted cultures were dropped onto the surface of swimming plates (0.3% agar starvation media) at sites equally distant from each other and from the sides of plates. After inoculation, the plates were incubated in a humidity-controlled environmental chamber at 30°C for 12 days, with daily images taken using a Panasonic Lumix GX85 Mirrorless Camera. Colony expansion was determined by measuring the lengths and width of the visible bacterial motility halos and calculating the average of the length and width for each colony.

### Statistical analyses

Mean values and standard errors (SEM) or median and quantiles with at least 3 technical replicates were calculated using GraphPad Prism 9.0. For the plant assay, box and whiskers plot were used to show the data distribution. Levels of significance were evaluated using either t-test or one-way ANOVA, followed by Tukey’s multiple comparison tests by GraphPad Prism 9.0. P-value large than 0.05 is flagged with non-significance (ns); P-value less than 0.05 is flagged with one star; P-value less than 0.01 is flagged with two stars; P-value less than 0.001 is flagged with three stars; P-value less than 0.0001 is flagged with four stars.

## Supporting information

SHC_Supplementary Tables

SHC_Supplementary Materials

## ACKNOWLEDGMENTS

We thank current and former members of the Belin lab for their helpful discussions. We are grateful to our colleagues in the Carnegie Embryology department for assistance with various aspects of the research, especially Dr. Mahmud Siddiqi for microscopy training and troubleshooting, and Dr. Frederick Tan, Dr. Alison Pinder, Dr. Xiaobin Zheng, and Dr. Karina Gutierrez for sequencing and related analysis. Dr. Birgit Scharf (Virginia Tech) and her laboratory gave indispensable advice and training for performing swimming motility assays in rhizobia. We also humbly acknowledge all members of our department’s Front Office, IT, and Facilities teams for making everything we do possible. This research was supported by funds from the Carnegie Endowment and National Institute for General Medical Sciences (NIGMS) grants R00GM126141 and R35GM147015 (both to B.J.B).

