## Supplementary material for "Hopanoid lipids promote soybean-*Bradyrhizobium* symbiosis": SHC_Supplementary Materials

**Content:**

**Supplementary tables (refer to zip files)**

Table S1. Primer sequences used for qRT-PCR

Table S2. Complete RNA-seq read counts of all treatments/replicates

Table S3: Relative expression of Pcu-*shc*+E compared to WT+E

Table S4. Up-regulated DGEs of Pcu-*shc*+E compared to WT+E

Table S5. Down-regulated DGEs of Pcu-*shc*+E compared to WT+E

Table S6. qRT-PCR information for verification of RNA-seq data

**Supplementary figures**

Figure S1. Growth curves ($OD_{600}$) of WT and Pcu-*shc* under varying cumate concentrations

Figure S2. Pcu-*shc* is an inefficient soybean symbiont at 24 dpi

Figure S3. Pcu-*shc* is an inefficient soybean symbiont at 45 dpi

Figure S4. Confocal sections of WT+E-infected soybean nodules at 27dpi

Figure S5. Confocal sections of low-occupancy soybean nodules infected with Pcu-*shc*+E at 27dpi

Figure S6. Confocal sections of high-occupancy soybean nodules infected with Pcu-*shc*+E at 27dpi

Figure S7. Pcu-*shc* fails to form symbioses with *Aeschynomene afraspera* at 28 dpi

Figure S8. Pearson correlation of WT and Pcu-*shc* with and without cumate

Figure S9. Top 20 up and down-regulated DEGs in Pcu-*shc*+E compared to WT+E

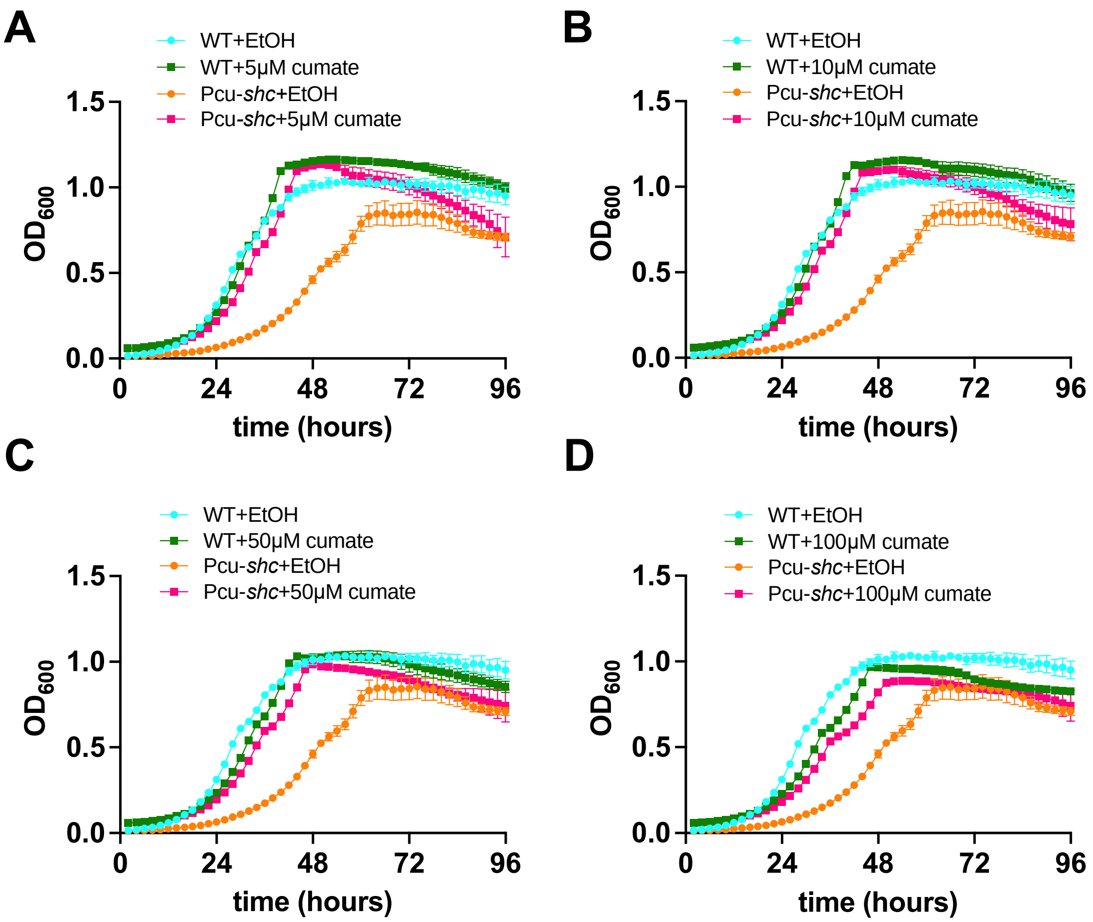


**Figure S1. Growth curves ($OD_{600}$) of WT and Pcu-*shc* under varying cumate concentrations.** *B.diazoefficiens* WT and Pcu-*shc* with and without cumate were grown for 4 days in AG media at pH of 6.6, 30˚C supplemented with EtOH only or with (**A**) 5 μM, (**B**)10 μM, (**C**) 50 μM, (**D**) 100 μM cumate

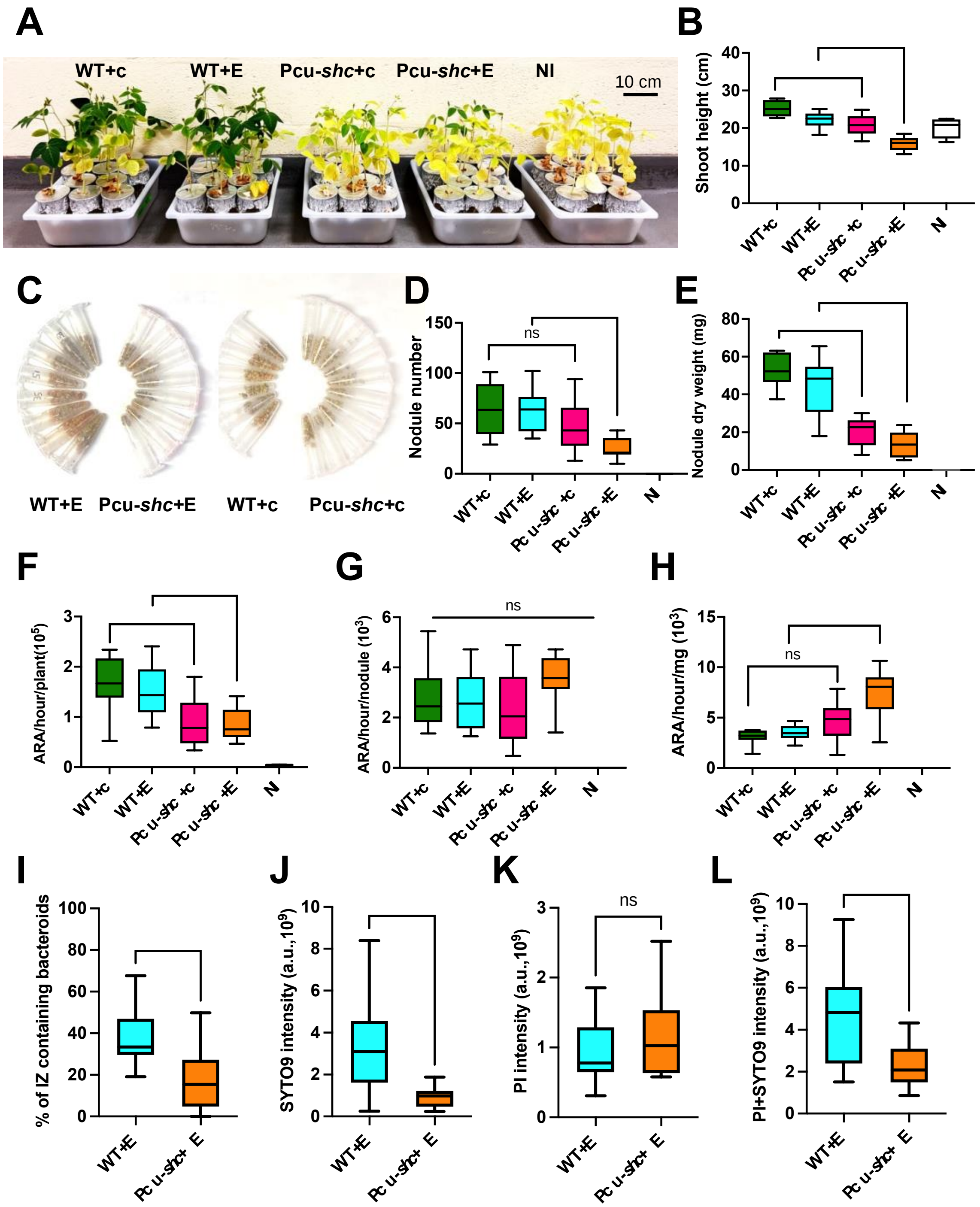

**Figure S2. Pcu-*shc* is an inefficient soybean symbiont at 24 dpi.**
**(A)** Comparison of growth of soybean inoculated with *B.diazoefficiens* WT and Pcu-*shc* with and without cumate. Non-inoculated (NI) soybeans are shown as controls. n=9 plants/treatment. **(B)** Median and quartiles of shoot height (cm) per plant in (A). **(C)** Images of nodules in 1.5 mL Eppendorf tubes collected from each plant in (A). (**D-E**) Median and quartiles of nodule number (C) and nodule dry weight (mg) per plant (D) for the treatments in (A). **(F-H)** GC-MS quantification of nitrogen fixation rate per plant by Acetylene Reduction Assay (ARA) normalized by reaction time (F), nodule number (G), and nodule weight (H). **(I-L)** ImageJ quantification of the portion of the infection zone (IZ) containing “LIVE” (SYTO 9) and “DEAD” (PI) bacteroids (I), and the intensity of “LIVE” (J), “DEAD” (K), and combined bacterial cells (L), a.u. = arbitrary units. Statistical analysis is described in Fig 3 and the Methods section.

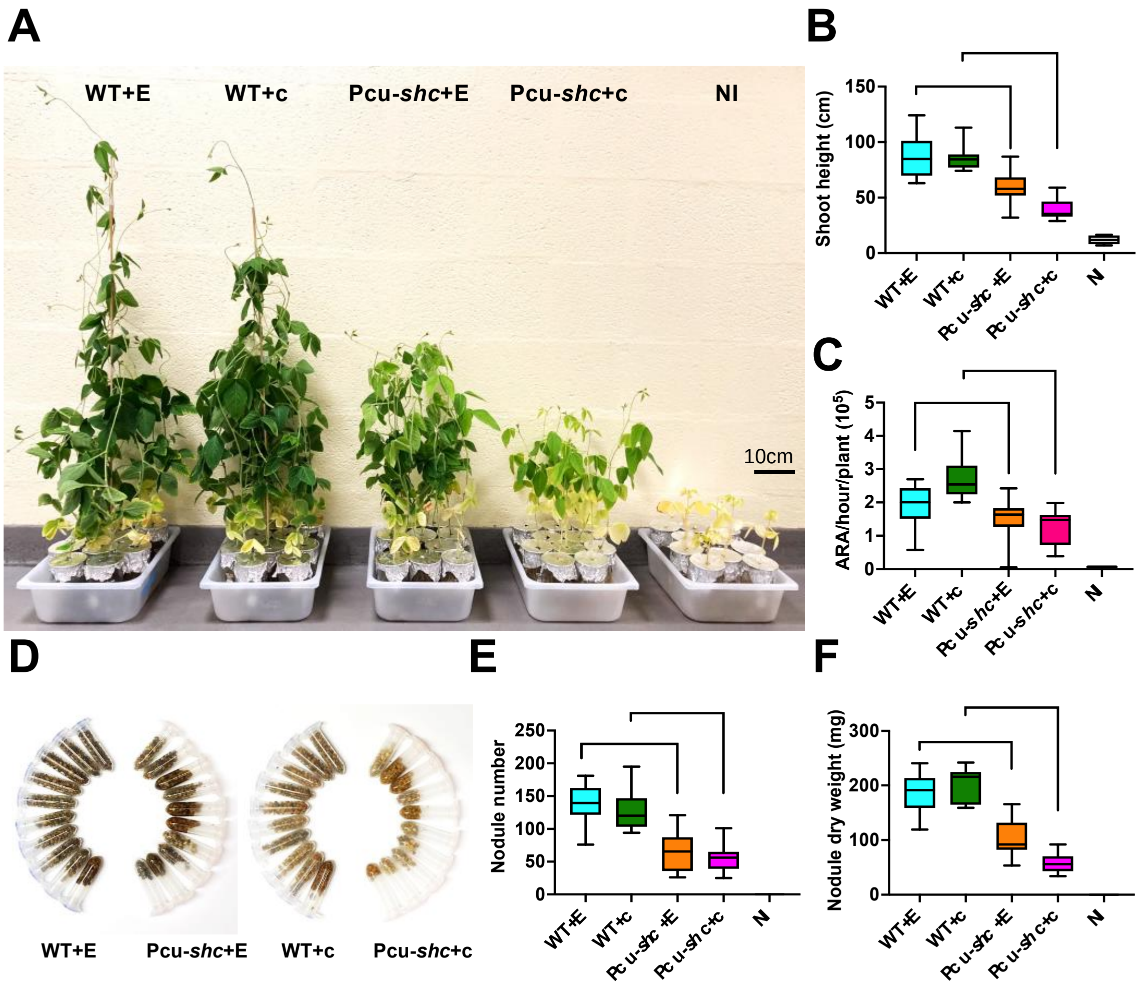


**Figure S3. Pcu-*shc* is an inefficient soybean symbiont at 45 dpi.**
**(A)** Comparison of growth of soybeans inoculated with *B.diazoefficiens* WT and Pcu-*shc* with and without cumate at 45 dpi. n=9 plants/treatment. **(B)** Median and quartiles of shoot height (cm) per plant in (A). **(C)** Median and quartiles of nitrogen fixation rate per plant quantified by GC-MS. **(D)** Images of nodules in 2 ml Eppendorf tubes collected from each plant in (A). (**E-F**) Median and quartiles of nodule number per plant and nodule dry weight (mg) per plant for the treatments of (A). Statistical analysis is described in Fig. 3 and the Methods section

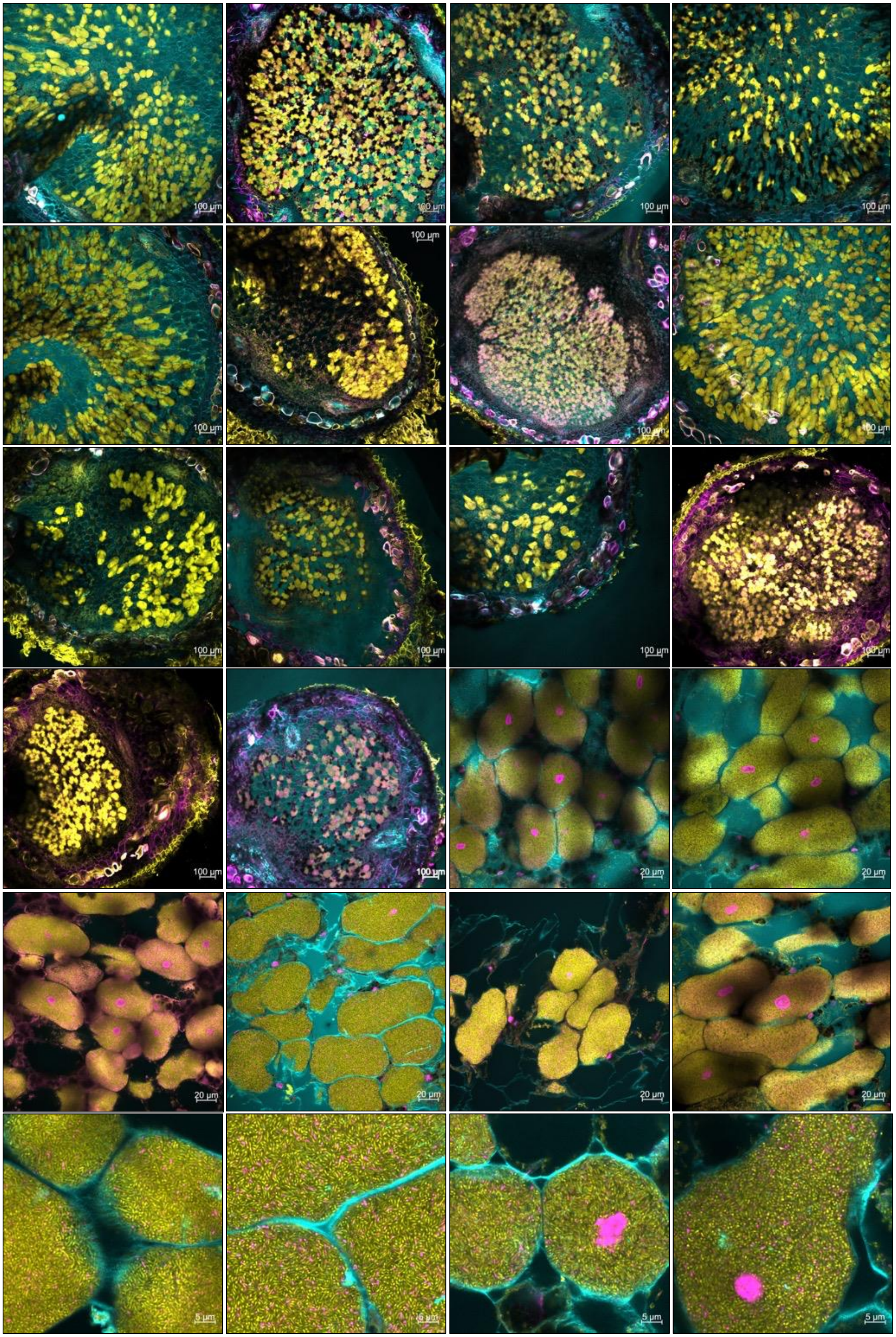


**Figure S4. Confocal sections of WT+E-infected soybean nodules at 27dpi.**
Nodule sections were stained with Calcofluor (cyan), SYTO 9 (yellow), and propidium iodide (magenta). Nodules were harvested from 5 plants.

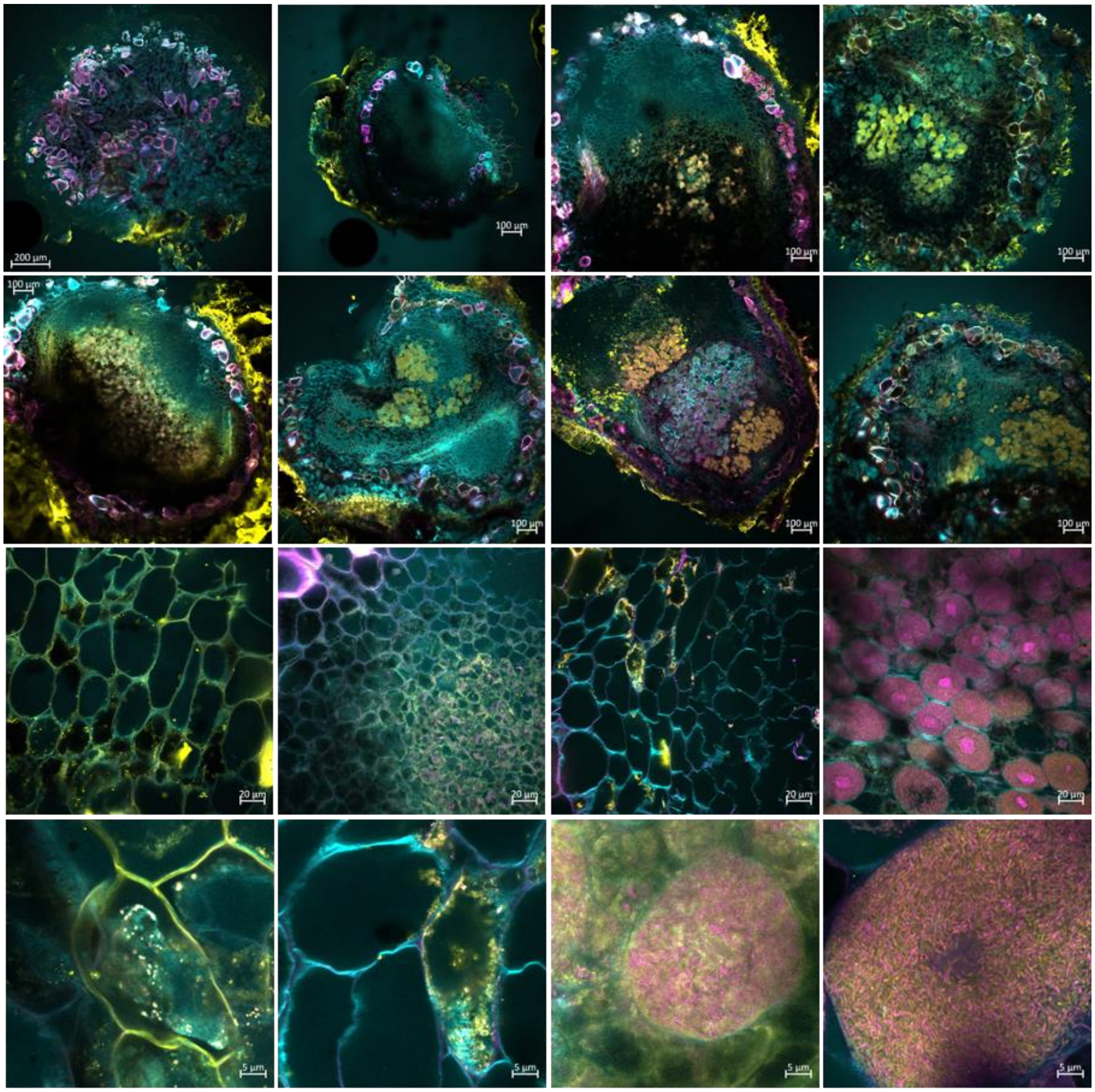

**Figure S5. Confocal sections of low-occupancy soybean nodules infected with Pcu-*shc*+E at 27dpi.** Nodule sections were stained with Calcofluor (cyan), SYTO 9 (yellow), and propidium iodide (magenta). Nodules were harvested from 5 plants.

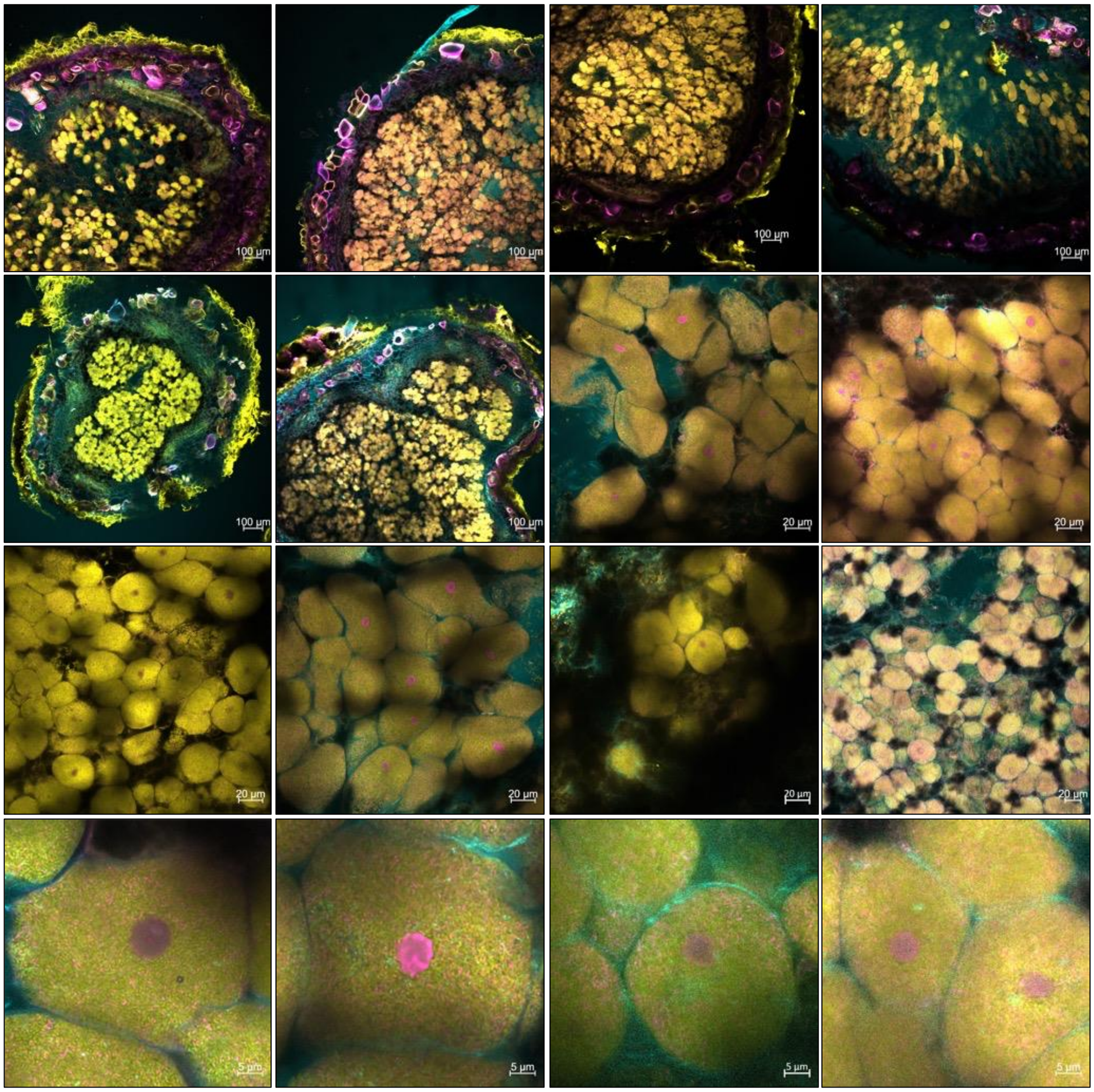


**Figure S6. Confocal sections of high-occupancy soybean nodules infected with Pcu-*shc*+E at 27dpi.** Nodule sections were stained with Calcofluor (cyan), SYTO 9 (yellow), and propidium iodide (magenta). Nodules were harvested from 5 plants.

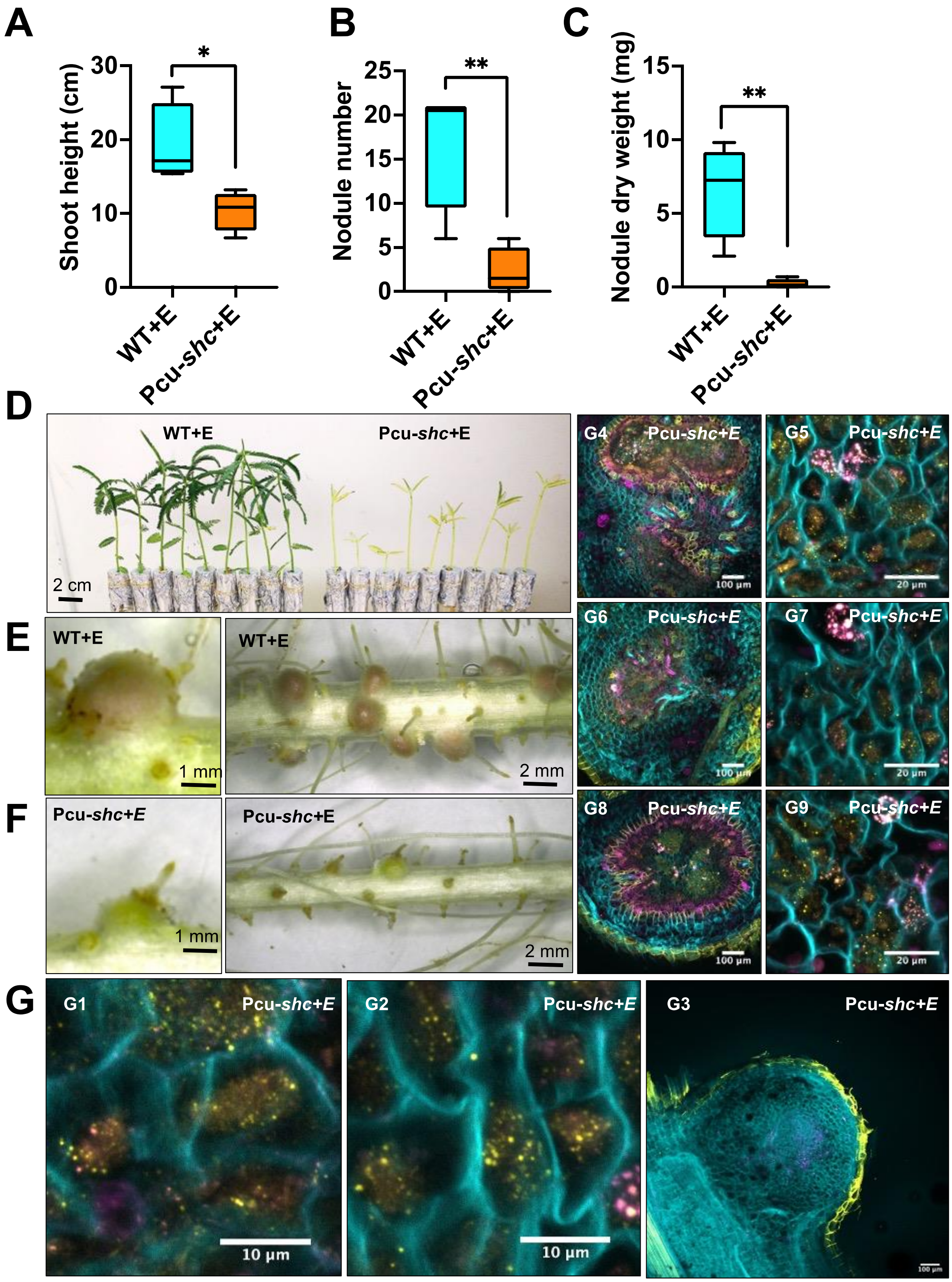


**Figure S7. Pcu-*shc* fails to form symbioses with *Aeschynomene afraspera* at 28 dpi.**
**(A-C)** Shoot height, nodule number, and nodule dry mass at 28 dpi for *A.afraspera* inoculated with *B.diazoefficiens* WT+E or Pcu-*shc*+E (9 plants/strain). **(D-F)** Images of whole plants and roots from inoculated WT+E or Pcu-shc+E plants. **(G)** Confocal images of nodule cross-sections from plants inoculated with Pcu-*shc*+E.

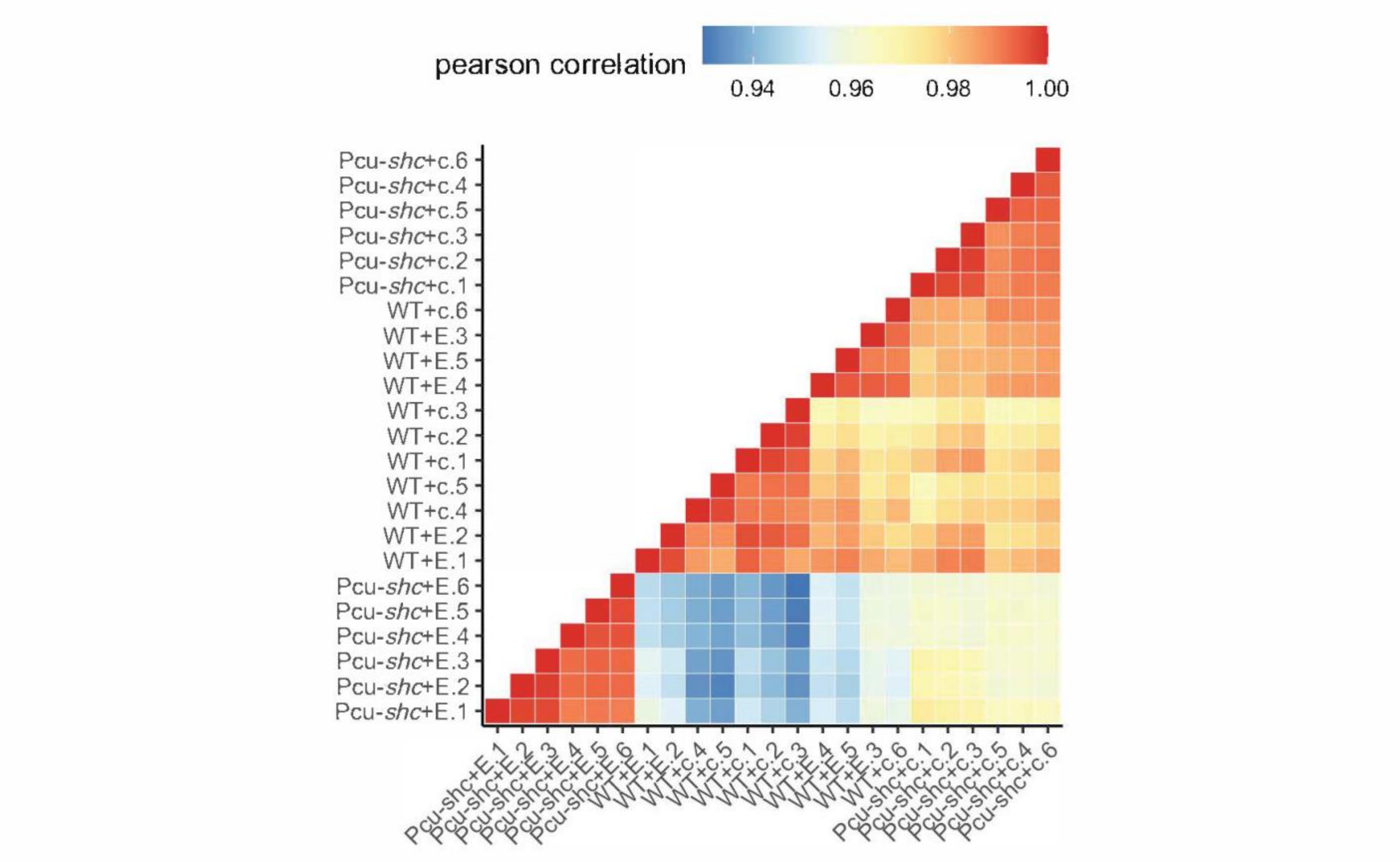


**Figure S8. Pearson correlation of WT and Pcu-*shc* with and without cumate.**
Pierson correlations of all RNA-seq replicates and treatments were analyzed by Deseq2 in Rstudio and were visualized by ImagGP (https://www.bic.ac.cn/ImageGP/)

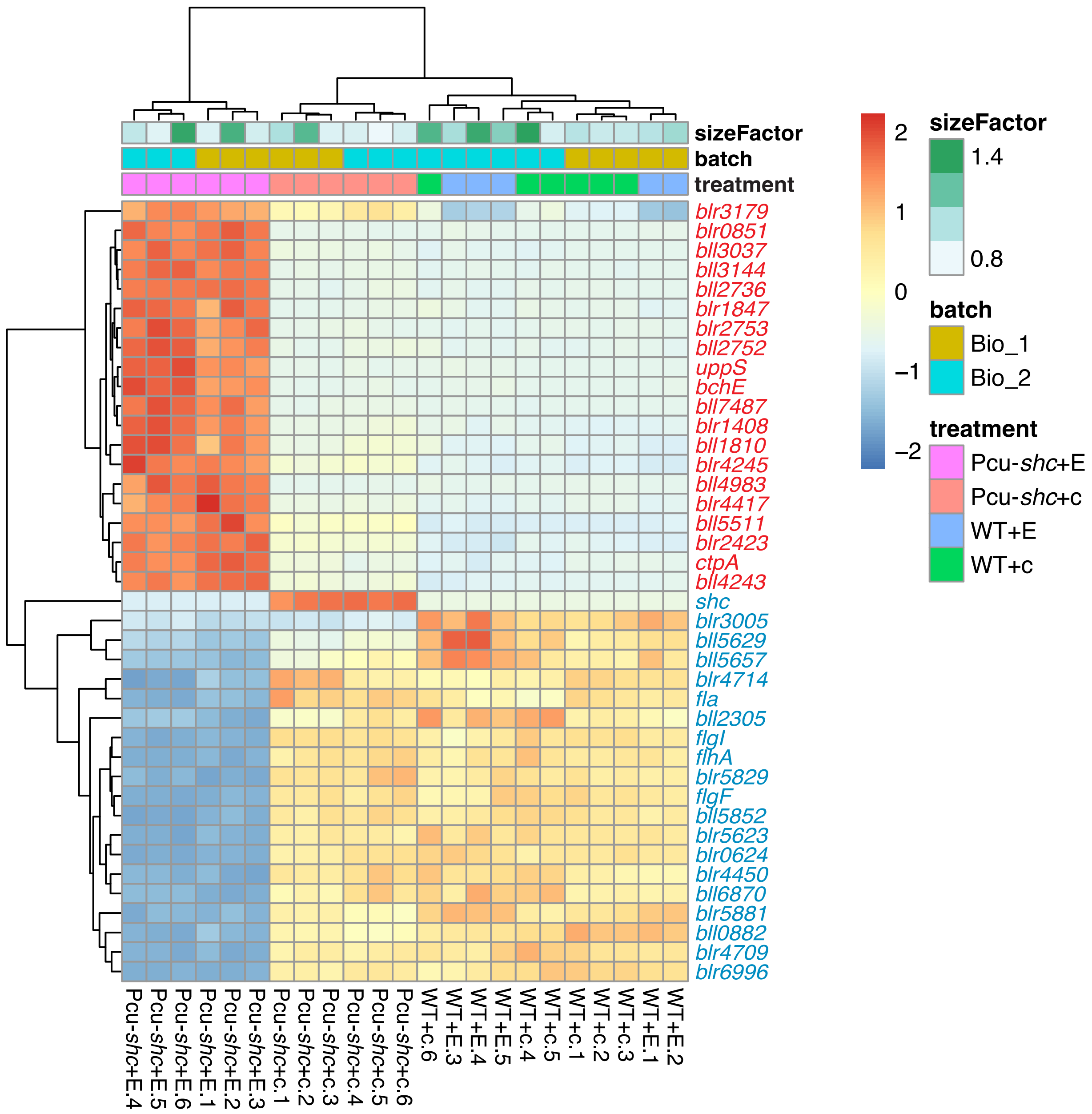


**Figure S9. Top 20 up and down-regulated DEGs in Pcu-*shc*+E compared to WT+E**.
Raw counts of WT and Pcu-*shc* with and without cumate from RNA-seq were normalized by the DESeq2's size factor and batch factor to correct for variability of the sequencing depth and batch effect from independent experiments. Heatmap was made by ImagGP from the top 20 up-and down-regulated DEGs of Pcu-*shc*+E compared to WT+E as shown in Table S4-S5, ranking by FDR. Bio_1 and Bio_2 represent two batches of samples collected from 09/2022 and 10/2022 respectively. Up-regulated genes are marked in red and down-regulate genes in blue.
